# Unraveling the host-selective toxic interaction of cassiicolin with lipid membranes and its cytotoxicity

**DOI:** 10.1101/2021.04.06.438527

**Authors:** Kien Xuan Ngo, Nguyen Bao Quoc, Phuong Doan N. Nguyen, Hirotoshi Furusho, Makoto Miyata, Tomomi Shimonaka, Nguyen Ngoc Bao Chau, Nguyen Phuong Vinh, Nguyen Anh Nghia, Tareg Omer Mohammed, Takehiko Ichikawa, Noriyuki Kodera, Hiroki Konno, Takeshi Fukuma, Toshio Ando

## Abstract

Cassiicolin (Cas), a toxin produced by *Corynespora cassiicola*, is responsible for corynespora leaf fall disease in rubber trees. Currently, the molecular mechanism of the cytotoxicity of Cas and its host selectivity have not been fully elucidated. Here, we analyzed the binding of Cas1 and Cas2 to membranes consisting of different plant lipids and their membrane-disruption activities. Using high-speed atomic force microscopy and confocal microscopy, we reveal that the binding and disruption activities of Cas1 and Cas2 on lipid membranes are strongly dependent on the specific plant lipids. The negative phospholipids, glycerolipids, and sterols are more susceptible to membrane damage caused by Cas1 and Cas2 than neutral phospholipids and betaine lipids. Cytotoxicity tests on rubber leaves of RRIV 1, RRIV 4, and PB 255 clones suggest that the toxins cause necrosis of rubber leaves, except for the strong resistance of PB 255 against Cas2. Cryo-SEM analyses of necrotic leaf tissues exposed to Cas1 confirm that cytoplasmic membranes are vulnerable to the toxin. Thus, the host selectivity of Cas toxin is attained by the lipid-dependent binding activity of Cas to the membrane, and the cytotoxicity of Cas arises from its ability to form biofilm-like structures and disrupt specific membranes.

**Graphic Abstract:** 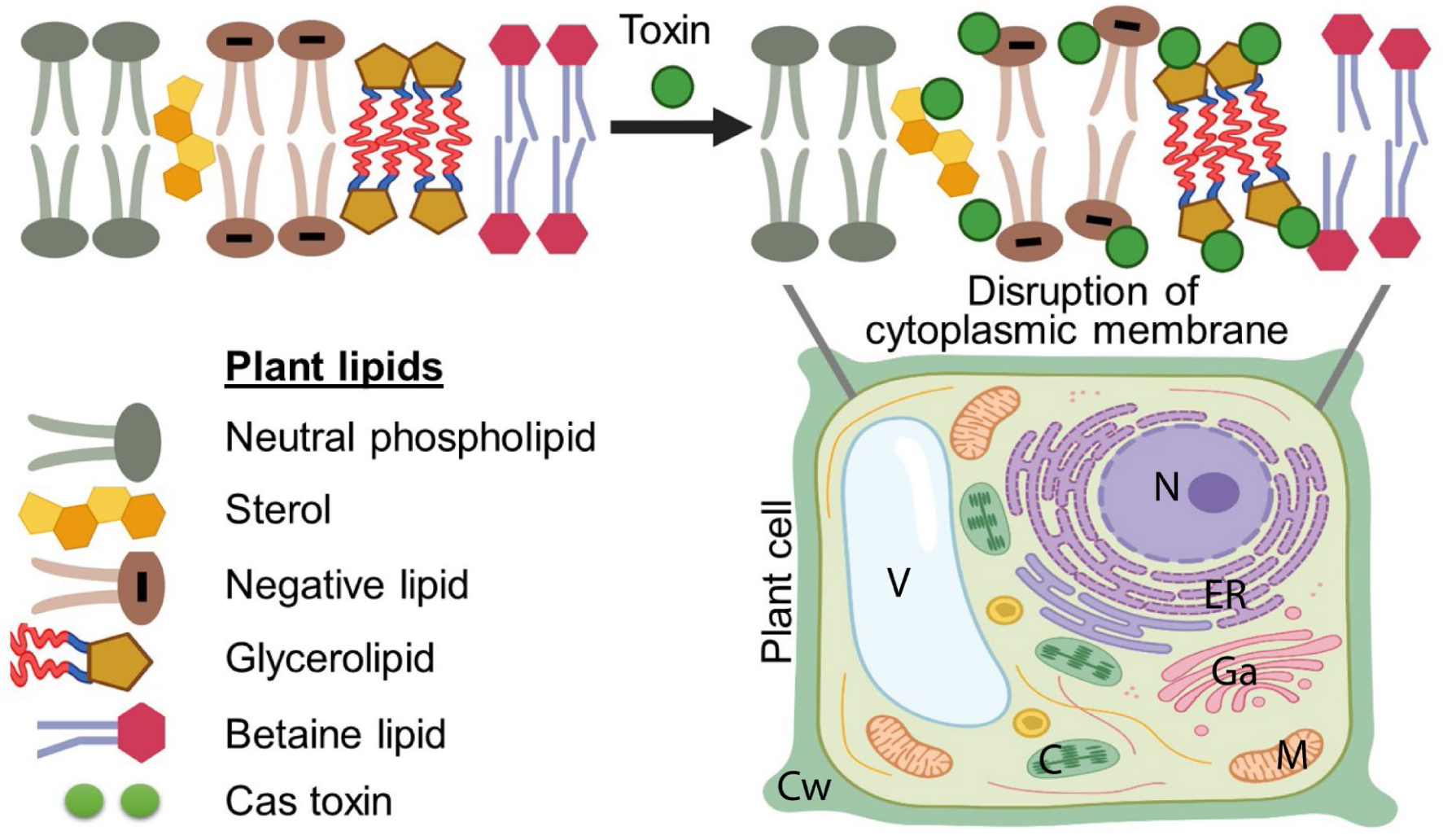

**Highlights:**

- Cas1 and Cas2 directly damage and cause necrosis in the leaves of specific rubber clones.
- Cas1 and Cas2 can form biofilm-like structures on specific lipid membranes (negative phospholipids, glycerolipids, and sterols). The biofilm-like formation of Cas toxin plays an important role in selective disruption on lipid membranes.
- Vulnerability of the specific cytoplasmic membranes to the selective Cas toxin that is the most remarkable feature of cytotoxicity of Cas toxin on plant cells.

## Introduction

The necrotrophic fungus *Corynespora cassiicola* (Berk.&Curt) Wei is a causal pathogen of corynespora leaf fall (CLF) disease in rubber leaves, of which the typical symptoms are brown leaf lesions surrounded by a yellow halo and occasional blackening of the veins, which causes a symptomatic “fish-bone” appearance, resulting in massive defoliation in susceptible rubber clones (Ribeiro et al., 2019). *C. cassiicola* predominantly infects both young and mature leaves, and CLF disease has now become widespread in *Hevea brasiliensis* plantations, causing severe damage to the natural rubber industry. These pathogens have also been found in many other crops (Dixon et al., 2009; Shrestha et al., 2017; Sumabat et al., 2018). Although the first pathogenic *C. cassiicola* isolate originally from the Philippines (CCP) is described as a causal agent in CLF disease, a necrotrophic effector known as cassiicolin (Cas) toxin produced in CCP is widely accepted as the host-selective toxin (HST) interaction in rubber trees and other plants (Barthe et al., 2007; de Lamotte et al., 2007; Déon et al., 2014, 2012a; Ribeiro et al., 2019; Wu et al., 2018). The Cas-encoding genes were found to be transiently expressed one or two days post inoculation, suggesting a role of Cas toxin in the early phase of infection (Déon et al., 2012a). Cas toxin is a small glycosylated secreted protein effector associated with secondary metabolism in *C. cassiicola* (Barthe et al., 2007; de Lamotte et al., 2007; Déon et al., 2014, 2012a). To date, six cassiicolin isoforms (Cas1 to Cas6) have been genetically identified by PCR-based detection from fungal isolates in various host plants and geographical origins (Déon et al., 2014, 2012a). Of note, *C. cassiicola* isolates only carry and express specific Cas genes, depending on the host specialization (Déon et al., 2014; Hieu et al., 2020; Lopez et al., 2018; Nghia et al., 2008; Nguyen et al., 2014). Some *C. cassiicola* isolates possessing no Cas gene yet causing CLF diseases may reveal other putative effectors involved (Lopez et al., 2018). The nature of the HST interaction of Cas toxin in CLF disease remains controversial because *in silico* analysis and prediction revealed 2870 putative effectors comprising CaZymes, lipases, peptidases, and secreted proteins and enzymes associated with secondary metabolism of *C. cassiicola* (Lopez et al., 2018). However, recent studies reported that gene deletion of Cas1 in *C. cassiicola* completely suppresses fungal pathogenicity in rubber trees (Ribeiro et al., 2019; Tran et al., 2016). These studies suggested two important findings: (i) Cas1 is a necrotrophic effector conferring virulence to *C. cassiicola* in susceptible rubber clones; (ii) other effectors produced in *C. cassiicola* contribute to residual filtrate toxicity and virulence in senescing/wounded tissues. Thus, the putative effectors may be involved in saprotrophs rather than necrotrophs. However, the cytotoxicity of Cas2 on rubber leaves has never been clarified before and remains unknown. Of special interest, we further examined and clarified the cytotoxicity of Cas isoforms (i.e., Cas1 and Cas2) to understand the fundamental molecular mechanisms underlying the HST interaction of Cas toxin in leaves of various rubber clones. In this context, the direct cytotoxicity of Cas1 to rubber leaves has been explored in some previous studies (Barthe et al., 2007; de Lamotte et al., 2007; Déon et al., 2012b). Notably, purified Cas1 applied on rubber leaves caused the same symptoms as *C. cassiicola* did at the cellular level (Barthe et al., 2007; de Lamotte et al., 2007). Electron microscopic analyses of necrotic leaf tissues revealed that Cas1 severely damaged plasma membranes in the chloroplast. Similarly, AK toxins from *Alternaria kikuchiana* and fumonisins and AAL toxin from *Fusarium* species infecting crops and plants were also reported to severely damage cytoplasmic membranes in various ways (Meena and Samal, 2019; Park et al., 1976; Riley et al., 1996; Tsuge et al., 2013). Objectively, no study has sufficiently clarified the susceptibility of plant lipids such as phospholipids, glycerolipids, sterols, and betaine lipids (Bae et al., 2020; Dörmann and Benning, 2002; Dufourc, 2008a, 2008b; Hartmann, 1998; Liengprayoon et al., 2011; Mackender and Leech, 1974; Murakami et al., 2018; Poincelot, 1976) to the pathogenicity of Cas toxin.

Alternatively, plant resistance to infectious diseases is also known to be associated with several defense responses activated by host-pathogen interactions (Thatcher et al., 2005). Recent transcriptomic analyses of the genetic responses of rubber clones susceptible and tolerant to *C. cassiicola* explored the diversity of genes encoding disease resistance proteins, leucine-rich repeat proteins, and genes involved in carbohydrate metabolic processes that were significantly upregulated in tolerant rubber clones upon infection. These genes, however, were either completely suppressed or downregulated in susceptible rubber clones (Roy et al., 2019). Again, molecular phylogenetic studies failed to determine the key factor in the host selectivity of Cas toxin. The nature of the HST interaction of Cas toxin in CLF disease in different rubber clones remains elusive.

This study addressed several key questions to elucidate the HST interaction of Cas toxin with plant lipids and its cytotoxicity on rubber leaves. Specifically, we examined the susceptibility of several important lipid components in plants to the pathogenicity of Cas1 and Cas2 isoforms at the molecular and cellular levels by combining high-speed atomic force microscopy (HS-AFM), confocal microscopy, and cryo-scanning electron microscopy (Cryo-SEM), thereby clarifying the HST interaction of Cas1 and Cas2 isoforms to different plant lipids. At the cellular level, we examined the cytotoxicity of Cas1 and Cas2 on rubber leaves of RRIV 1, RRIV 4, and PB 255 clones to understand the HST interaction of Cas isoforms to specific hosts. Here, we report that the binding and disruption activities of Cas1 and Cas2 are more sensitive to negatively charged lipids (DPPA), glycerolipids (MGDG, DGDG), and sterols (stigmasterol, sitosterol) than to neutral (DPPC) and betaine lipids (DGTS-d9, DGTS). Cas toxins could form biofilm-like structures onto specific lipid membranes composed of negative phospholipid, glycerolipids, and sterols, but failed to form these structures onto the membranes composed of betaine lipids (e.g., DGTS-d9). Cytotoxicity tests of Cas1 and Cas2 on rubber leaves of RRIV 1, RRIV 4, and PB 255 clones suggest that these clones are highly susceptible to the pathogenicity of Cas1 and Cas2 except for the strong resistance of PB 255 against Cas2. Finally, we also report that cytoplasmic membranes in susceptible rubber clones are highly vulnerable to the pathogenicity of purified Cas toxin and toxin produced from fungus *C. cassiicola*.

## Results

### Disruption activity of Cas1 and Cas2 to different lipid membranes imaged by HS-AFM

To examine the possible pathogenicity of Cas1 and Cas2, this study addressed the first question of whether Cas1 and Cas2 could bind to and disrupt various phospholipids (DPPC, DPPA), glycerolipids (MGDG, DGDG), sterols (stigmasterol, sitosterol), and betaine lipids (DGTS-d9, DGTS) available in plants. To answer this question, we used tapping-mode HS-AFM to image the dynamic interaction and disruption activities of Cas1 and Cas2 to different lipid membranes in real time. Unlike a conventional AFM, which has a slow scanning rate, HS-AFM can image in sub-second temporal resolution many biological molecules during functional activities (Ando, 2012; Ando et al., 2013, 2008, 2007; Uchihashi et al., 2012). The very soft and small cantilever used in HS-AFM was proven in numerous previous reports to non-invasively perturb the structures and functions of biological molecules (Kodera et al., 2010; Lin et al., 2020; Ngo et al., 2015; Uchihashi et al., 2011). This microscopy enables real-time, high-resolution imaging of individual biological nanomachines during their functional activities, as demonstrated by studies on walking myosin V (Kodera et al., 2010), rotorless F1-ATPase (Uchihashi et al., 2011), and actin filaments interacting with cofilin (Ngo et al., 2015), and recently explores many undiscovered features of many other inspired biomolecules (Ando, 2018; Heath and Scheuring, 2019; Karner et al., 2017; Kodera et al., 2020; Lin et al., 2020; Nasrallah et al., 2019; Shibata et al., 2017).

We produced Cas1 and Cas2 recombinant proteins in *E. coli*. The basic characteristics of these proteins are described in **Figure 1-figure supplements 1, 2, and Figure 1-tables 1, 2**. Our analysis predicted that a signal peptide (SP) domain in Cas1 and Cas2 containing several different hydrophobic amino acids (M, Y, L, I, A, F, V) may help to better insert these proteins into the hydrophobic region of a lipid bilayer. The numbers of negatively and positively charged residues in Cas1 and Cas2 are identical. However, the secondary structures (α-helix, β-sheet, and β-turn) in the mature domains of Cas1 and Cas2 are locally different (see red and yellow rectangles in **Figure 1-figure supplement 1**). To follow the binding and disruption activities of Cas1 and Cas2 on preformed lipid membranes in solution, Cas1 or Cas2 was added into the imaging chamber at a final concentration of 0.95 µM during HS-AFM imaging. Initially, we found that neutral phospholipid (DPPC) membranes were weakly sensitive to the binding and disruption activities of Cas1 and Cas2 (**Figures 1a1, a2, d-a1, d-a2, Figure 1-figure supplement 3, Table 1, and Videos 1 – 2**). Thus, we made several composite lipid membranes by mixing with DPPC lipids to investigate their susceptibility to the pathogenicity of Cas1 and Cas2. As shown in **Figures 1b1, d-b1, c1, d-c1, d1, d-d1, b2, d-b2, c2, d-c2, d2, and d-d2, Table 1, and Videos 3 – 8**, the negatively charged phospholipid (DPPC/DPPA) and glycerolipid (DPPC/MGDG, DPPC/DGDG) mixtures were highly sensitive to the binding and disruption activities of Cas1 and Cas2. The DPPC/stigmasterol and DPPC/sitosterol membranes were also strongly susceptible to the binding and disruption activities of both Cas1 and Cas2 (**Figures 1e1, d-e1, e2, d-e2, f1, d-f1, f2, and d-f2, Table 1, and Videos 9 – 12**). The lipid membrane mixtures of betaine lipids (DPPC/DGTS-d9, DPPC/DGTS) were weakly vulnerable to Cas1 and Cas2 (**Figures 1g1, d-g1, h1, d-h1,** g2, d-g2, h2, and d-h2, Table 1, and Videos 13 – 16).

**Figure 1.**
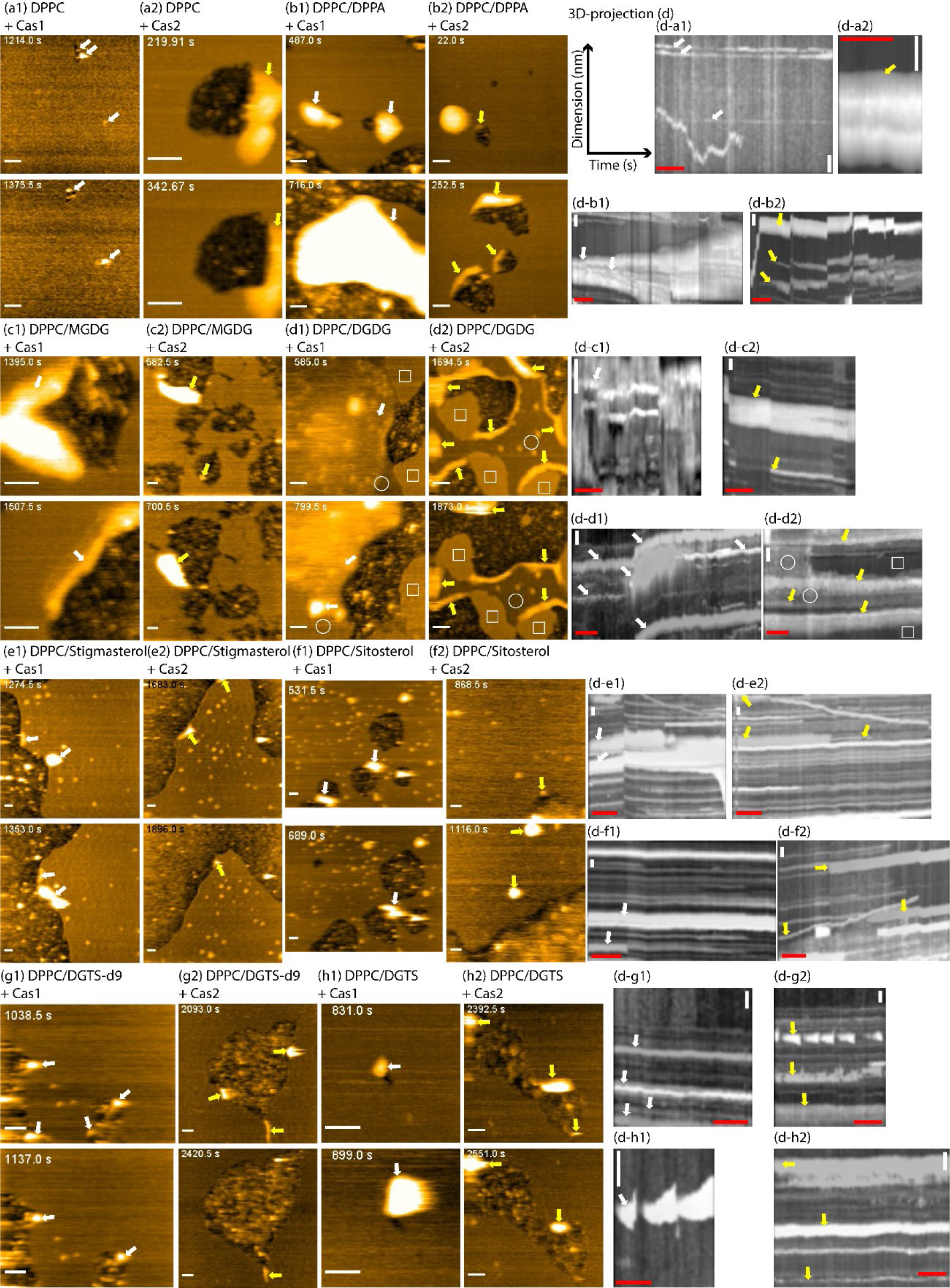
HS-AFM imaging of disruption processes of different membrane lipids available in plants induced by Cas1 and Cas2. **d-a1 – d-h2**: 3D projection analyses corresponding to a1 – h2 and Videos 1 – 16, respectively, to track the dynamics of Cas clusters formed and incorporated onto lipid membranes. Briefly, lipid bilayers consisting of different lipid compositions were formed on mica by rupturing liposomes. Then, 0.95 µM Cas1 or Cas2 was added, and HS-AFM imaging was carried out in C-buffer using a BL-AC10DS-A2 cantilever deposited with a small lab-made tip. White and yellow arrows: Cas1 and Cas2 clusters, respectively, incorporated into the lipid bilayer. Rectangles and circles in d1 and d2 denote DPPC and DGDG domains, respectively. Imaging rate: 2 fps. The time label in still images denotes a time point after the addition of Cas1 or Cas2. In kymographs, white bars: 50 nm, red bars: 25 s. **<u>Related to Videos 1</u>** <u>– 16, and</u> Table 1.

**Table 1.**
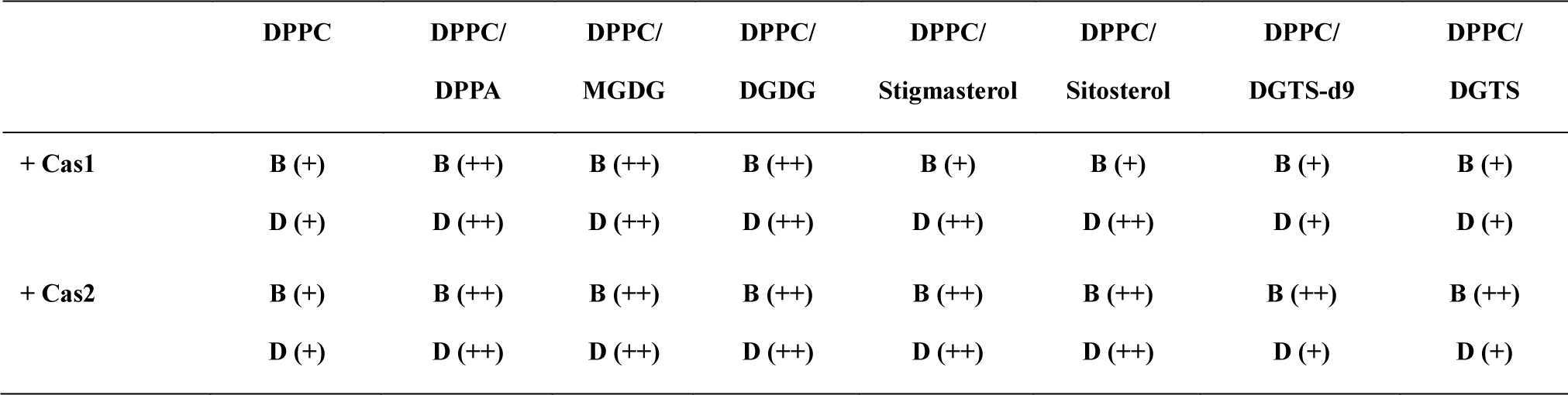
Classification of the susceptibility of different lipids available in plant cytoplasmic membranes to Cas1 and Cas2. Here, we compared the binding and disruption activities of Cas1 and Cas2 onto the lipid membranes acquired in HS-AFM and confocal microscopic analyses. The binding and disruption activities of Cas1 and Cas2 to the neutral DPPC lipid membranes were used as the basal indicator for a semi-quantitative comparison. **++**: strong; **+**: weak; B: Binding; D: Disruption. **<u>Related to</u> Figures 1<u>, 2, 3, and 4, and Videos 1 – 16</u>**.

In this study, our real-time HS-AFM imaging Videos revealed that Cas1 and Cas2 were strongly bound to membranes containing negatively charged phospholipid (DPPA), glycerolipids (MGDG, DGDG), and sterols (stigmasterol, sitosterol) and penetrated deeply into the lipid membranes. Cas1 and Cas2 formed large clusters on these highly susceptible membrane lipids and expanded their clusters, similar to tsunamis, to devastate the membranes (**Figures 1b1, b2, c1, c2, d1, d2, e1, e2, f1, and f2, and Videos 3 – 10**). By acting in these ways, they continuously damage lipid membranes. After disruption, many aggregates left on mica were observed. These aggregates were likely a mixture composed of lipids and Cas1 or Cas2. Our results also show that neutral DPPC lipids were weakly susceptible to the binding and disruption activity of both Cas1 and Cas2 (**Figures 1a1, d-a1, a2, d-a2, and Videos 1 – 2**). This statement is firmly supported in the observation of the disruption activity of Cas1 and Cas2 to DPPC domains and DGDG domains segregated in DPPC/DGDG membranes (**Figure 2B, 2D, and Videos 7 – 8**). The DPPC and DGDG lipid domains formed on the mica surface were distinguished by a height difference of ∼1.5 – 1.9 nm. Cas clusters were also distinguished by measuring the height differences with membranes and are representatively shown for Cas1 and Cas2 clusters (**Figure 2**). DGDG domains were strongly devastated by Cas1 and Cas2 clusters, while DPPC domains remained almost intact (kymographs in **Figures 2E and 2F**). Based on this observation, we further confirmed that glycerolipids (MGDG, DGDG) are strongly susceptible to the disruption activity of Cas1 and Cas2. Most likely, the negatively charged surface and high membrane fluidity of glycerolipids are two key factors for better binding, accessibility, and damage activity of Cas1 and Cas2 clusters in these membranes. We further tested membrane accessibility for the disruption activity of Cas1 and Cas2 by incorporating sterols (stigmasterol, sitosterol) into the membranes. The presence of plant sterols in membranes was expected to increase the membrane accessibility for Cas1 and Cas2, although sterols are generally thought to decrease the membrane fluidity (Dufourc, 2008a, 2008b). Indeed, these membranes were greatly devastated by Cas1 and Cas2. The disruption activities of Cas toxin on these membranes happened regardless of the Cas cluster formation or not (**Figures 1e1, d-e1, e2, d-e2, f1, d-f1, f2, and d-f2 and Videos 9 – 12**). Additionally, when neutral betaine lipids (DGTS- d9, DGTS) were mixed with DPPC lipids, these membranes were only weakly damaged by Cas1 and Cas2. The degrees of damage of Cas1 and Cas2 on these membranes were similar to those on pure DPPC membranes (**Figures 1a1, d-a1, a2, d-a2, g1, d-g1, g2, d- g2, h1, d-h1, h2, and d-h2, and Videos 1 – 2, 13 – 16)**. Overall, we suggest that negatively charged phospholipids, glycerolipids, and specific sterols available in the cytoplasmic membranes of plant cells are strongly sensitive to the disruption activity of Cas1 and Cas2. Neutral phospholipids and betaine membrane lipids are weakly susceptible to the disruption activity of Cas1 and Cas2.

**Figure 2.**
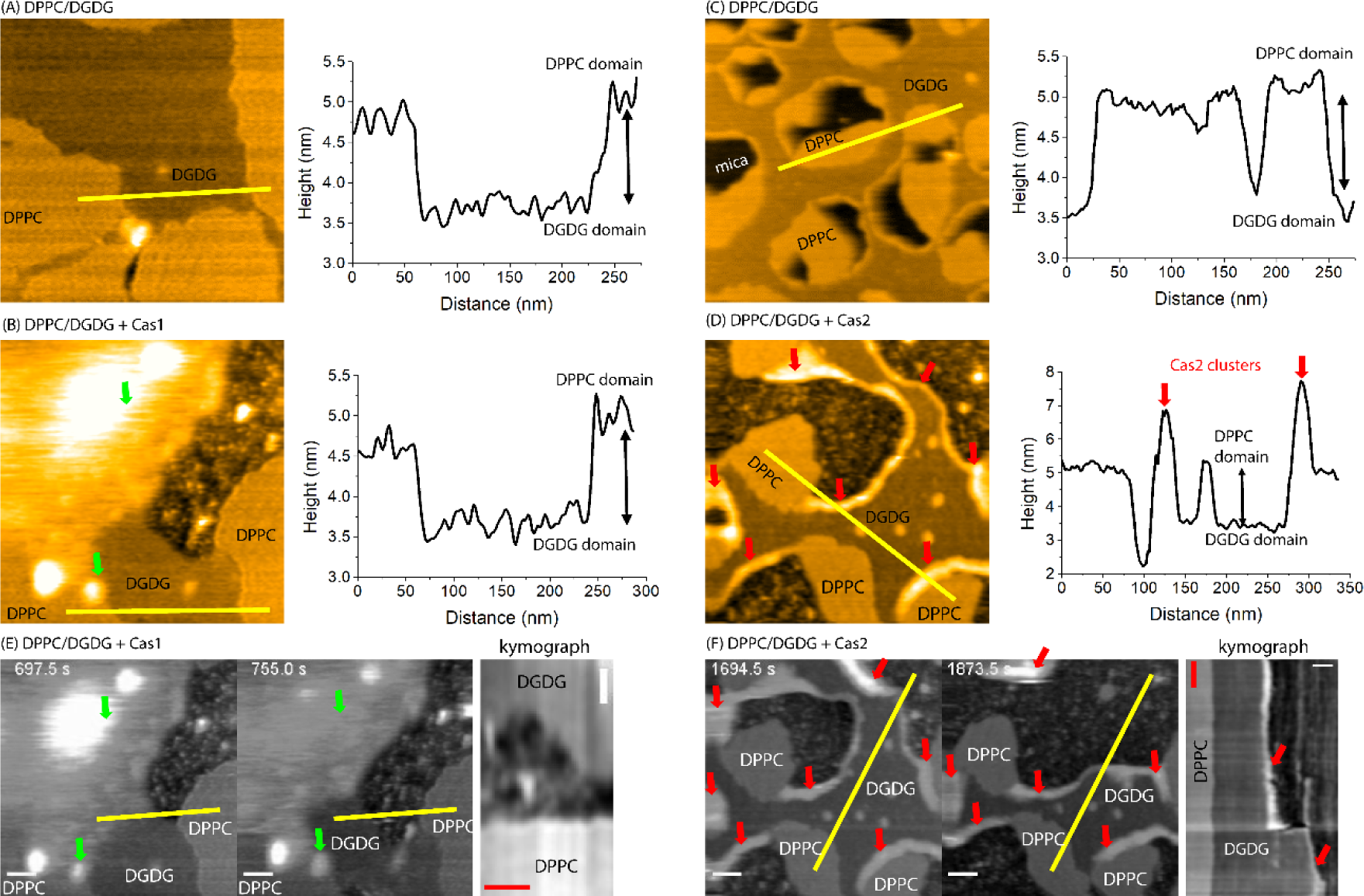
HS-AFM topographic image analyses. (A, B) HS-AFM images and topographic profiles of the DPPC/DGDG lipid bilayer formed on mica in the absence and presence of Cas1. (C, D) HS-AFM images and topographic profiles of the DPPC/DGDG lipid bilayer formed on mica in the absence and presence of Cas2. (E, F) Kymographs demonstrate the selective disruption activities of Cas1 and Cas2, respectively, on DPPC and DGDG membrane domains. DPPC, DGDG, and Cas clusters were distinguished by height differences. The time label in still images denotes a time point after the addition of Cas1 or Cas2. Green arrows: Cas1 clusters, red arrows: Cas2 clusters. In kymographs, white bars: 50 nm, red bars: 25 s. **<u>Related to Figures 1d1 – d2 and Videos 7 – 8.</u>**

### Biofilm-like formation of Cas toxin onto specific lipid membranes

To examine whether biofilm-like structures of Cas toxin could be formed on specific lipid membranes, we used HS-AFM and imaged in realtime the binding of Cas toxin to the preformed lipid membranes in solution. Generally, the formation of the infectious microbial biofilms plays various important roles such as tolerance to the host immune system, cell adhesion, serve as a reservoir for cell division (Mancera et al., 2020), and nutritive absorption, protective barrier, development of reproductive and vegetative structures (i.e., fungal spore and hyphae), and sexual and asexual reproduction (Gour et al., 2014; Siqueira and Lima, 2013).

In this study, we used a high concentration of Cas1 and Cas2 (4.75 µM) and compared the biofilm-like formation of Cas toxin on two different lipid membrane surfaces (i.e., the susceptible DPPC/sitosterol vs. the non-susceptible DPPC/DGTS-d9 membranes). We found that the lipid membranes composed of DPPC/sitosterol strongly recruited the binding and biofilm-like formation of the Cas1/Cas2 mixture (**Figure 3A, Video 17**). The biofilm of the Cas1/Cas2 mixture was covered and partially incorporated onto membranes. The biofilm thickness of the Cas1/Cas2 mixture was approximately 5 – 6.5 nm, and its height protruded over the membrane surface was ∼1 – 1.5 nm (**Figure 3A**). Consequently, the disruption activities of the Cas1/Cas2 mixture on these membranes were very strong (**Figure 3A, Video 17**). These membranes also recruited the biofilm-like formation of Cas2. The biofilm-like structures of Cas2 were covered on the membrane surface with a thickness of ∼0.3 – 0.5 nm (**Figure 3B, Video 18).** Contrarily, the lipid membranes made of DPPC/DGTS-d9 failed to form the biofilm-like structures of the Cas1/Cas2 mixture. Despite a combination of Cas1 and Cas2 at a high concentration, the Cas1/Cas2 clusters were partially incorporated into membranes and yet weakly disrupted these membranes (**Figure 3C, Video 19**). Additionally, Cas1 and Cas2 at a lower concentration (∼0.95 µM) could also form biofilm-like structures on membranes containing negative phospholipid (DPPC/DPPA) (**Figures 1b1, b2, Videos 3 – 4**) and glycerolipid (DPPC/DGDG) (**Figures 1d1, d2, Videos 7 – 8**). Biofilm-like formation of Cas1 was also seen on membranes containing glycerolipid (DPPC/MGDG) (**Figure 1c1, Video 5**). We suggest here, for the first time, that biofilm-like structures of Cas toxin can be formed onto specific lipid membranes available in plant cells (i.e., negative phospholipid, glycerolipids, and sterol). The effective biofilm-like formation of the Cas1/Cas2 mixtures onto specific lipid membranes may also reflect the nature of Cas toxin production and virulence of *C. cassiicola* possessing various Cas isoforms when they infect and grow on rubber leaves.

**Figure 3.**
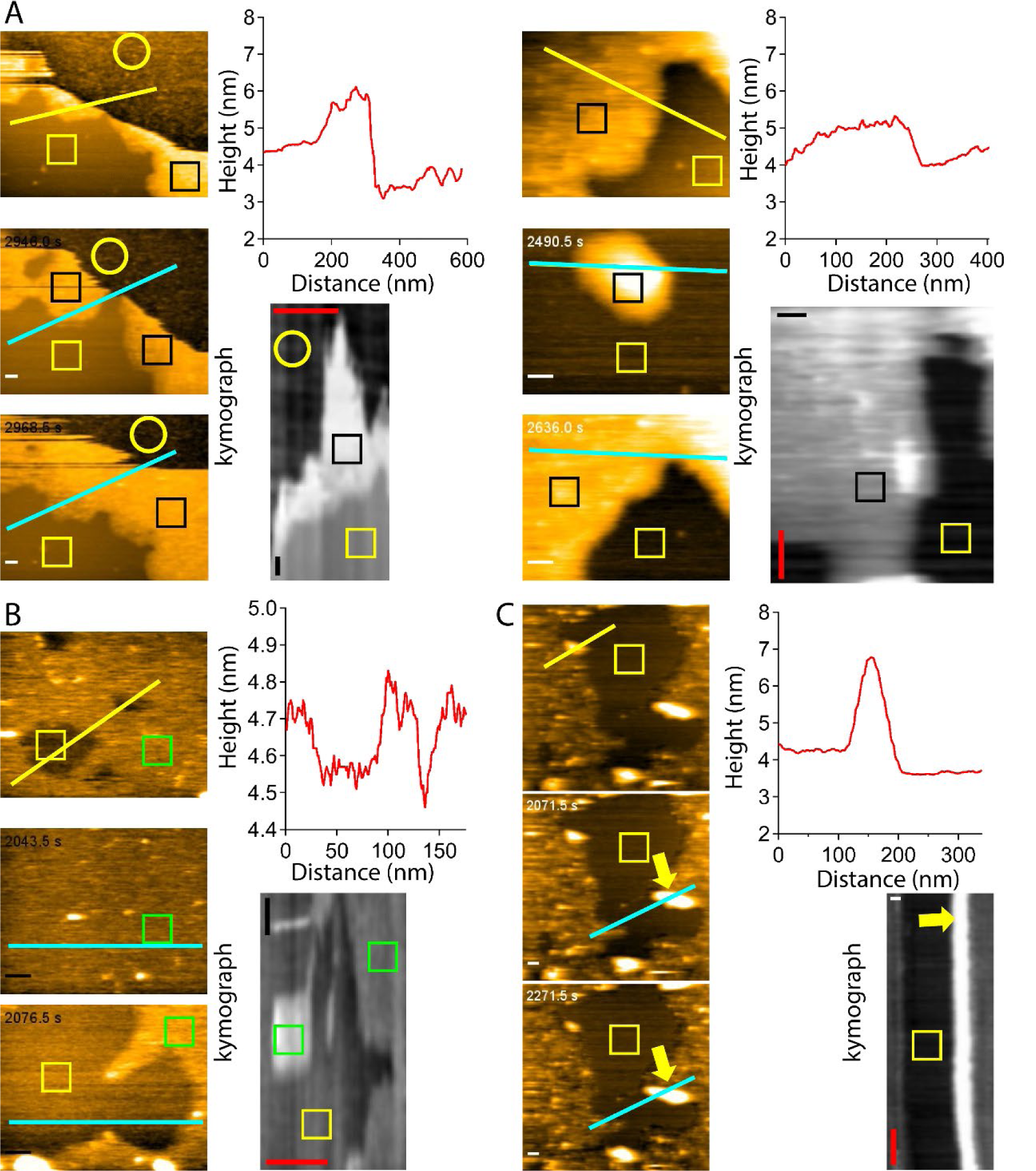
Biofilm-like structures of the Cas1/Cas2 mixture formed on the specific lipid membranes. (A) Biofilm-like structures of the Cas1/Cas2 mixtures formed on DPPC/sitosterol (80/20 wt%) membranes; (B) Biofilm-like structures of Cas2 formed on DPPC/sitosterol (80/20 wt%) membranes; (C) No formation of the biofilm-like structures of the Cas1/Cas2 mixtures on DGTS-d9 (80/20 wt%) membranes. The final concentration of Cas1 and Cas2 were 4.75 µM. The topographic profiles of biofilm-like structures are shown (yellow and red lines). The kymographs demonstrate the dynamics of biofilm-like formation as a function of time (cyan lines). The time label in still images denotes a time point after the addition of Cas1 and Cas2. Yellow arrows denote Cas clusters formed on DPPC/DGTS-d9 membranes. Black, yellow, and green rectangles denote biofilm-like areas (Cas1/Cas2 mixtures), lipid membranes, and biofilm-like areas (Cas2), respectively. Yellow circles denote the lipid/Cas1/Cas2 mixtures left on mica after the disruption process. In kymographs, white and black bars: 50 nm, red bars: 25 s. **<u>Related to Videos 17, 18, and 19</u>**.

### Binding of Cas1 and Cas2 to artificial cytoplasmic membranes imaged by confocal microscopy

We further examined the susceptibility of intact lipid membranes to the binding of Cas1 and Cas2 using confocal microscopy. Thus, we fabricated giant liposome vesicles (GVs) and used them as a model of artificial cytoplasmic membranes to assay the binding activities of GFP-Cas1 and GFP-Cas2 using confocal microscopy, a powerful technique to study protein-lipid membrane interaction (Carvalho et al., 2008; Drücker et al., 2019). The composite GVs composed of similar lipid compositions used in HS-AFM observation were made by electroformation methods (**Figure 1-figure supplement 3 and Figure 4-table 1**). Here, we mainly compared the binding of GFP-Cas1 and GFP-Cas2 to different GVs made of DPPC, DPPC/DPPA, DPPC/MGDG, DPPC/DGDG, DPPC/stigmasterol, DPPC/sitosterol, DPPC/DGTS-d9, and DPPC/DGTS lipid mixtures with the small addition of rhodamine-PE and biotinyl-PE lipids to absorb and image GVs. Principally, we fabricated GVs containing both giant unilamellar and multilamellar vesicles and used them to assay the binding activity of GFP-Cas1 and GFP-Cas2 (**Figure 4, Tables 1, 2**). The binding of GFP-Cas1 and GFP-Cas2 to GVs made of only neutral phospholipids (DPPCs) was indeed weak (**Figures 4a1, a2, i1, i2**). Most DPPC GVs were not bound with GFP-Cas1 and GFP-Cas2. Thus, we used DPPC as the basal lipid membrane to compare the binding degrees of GFP-Cas1 and GFP-Cas2 to GVs composed of negative phospholipids (DPPC/DPPA), glycerolipids (DPPC/MGDG, DPPC/DGDG), sterols (DPPC/stigmasterol, DPPC/sitosterol), and neutral betaine lipids (DPPC/DGTS- d9, DPPC/DGTS). GFP-Cas1 was bound more strongly to GVs composed of DPPC/DPPA, DPPC/MGDG, and DPPC/DGDG lipids than to GVs made of only DPPC (**Figures 4a1, b1, c1, d1, i1**). The susceptibility of GVs composed of DPPC/stigmasterol, DPPC/sitosterol, DPPC/DGTS-d9, and DPPC/DGTS lipids to the binding of GFP-Cas1 was also recorded as strong (**Figures 4e1, f1, g1, h1**), although when compared to the binding of GFP-Cas1 to GVs made of only DPPC (**Figure a1**), the difference (p ≤ 0.05, *t-test*) was not significant (**Figure 4i1, Table 2**). Additionally, we found that GFP-Cas2 bound more strongly to all GVs composed of the negative phospholipids, glycerolipids, sterols, and betaine lipids used in this study compared to GVs made of only the neutral DPPC phospholipids (**Figures 4a2, b2, c2, d2, e2, f2, g2, h2, i2**). These results strongly support that both GFP-Cas1 and GFP-Cas2 were bound to suspended lipid membranes, similar to artificial cytoplasmic membranes composed of the specific plant lipids. The binding of GFP-Cas1 and GFP-Cas2 was varied and dependent on the types of lipids. Although we did not evaluate the disruption of these lipid membranes caused by GFP- Cas1 and GFP-Cas2 using confocal microscopy, we consistently captured the large bright clusters of GFP-Cas1 and GFP-Cas2 formed in the susceptible lipid membranes (see GFP fluorescence in **Figures 4b1, b2, c1, d1, e2, f2, g2, h2**). This observation also implies that the susceptible lipid components available in plants (PA, MGDG, DGDG, stigmasterol, and sitosterol) recruited the binding and formation of Cas1 and Cas2 clusters in the lipid membranes, as seen in our HS-AFM observation. Most likely, the formation and expansion of the large Cas1 and Cas2 clusters in susceptible lipid membranes play crucial roles in devastating lipid membranes.

**Figure 4.**
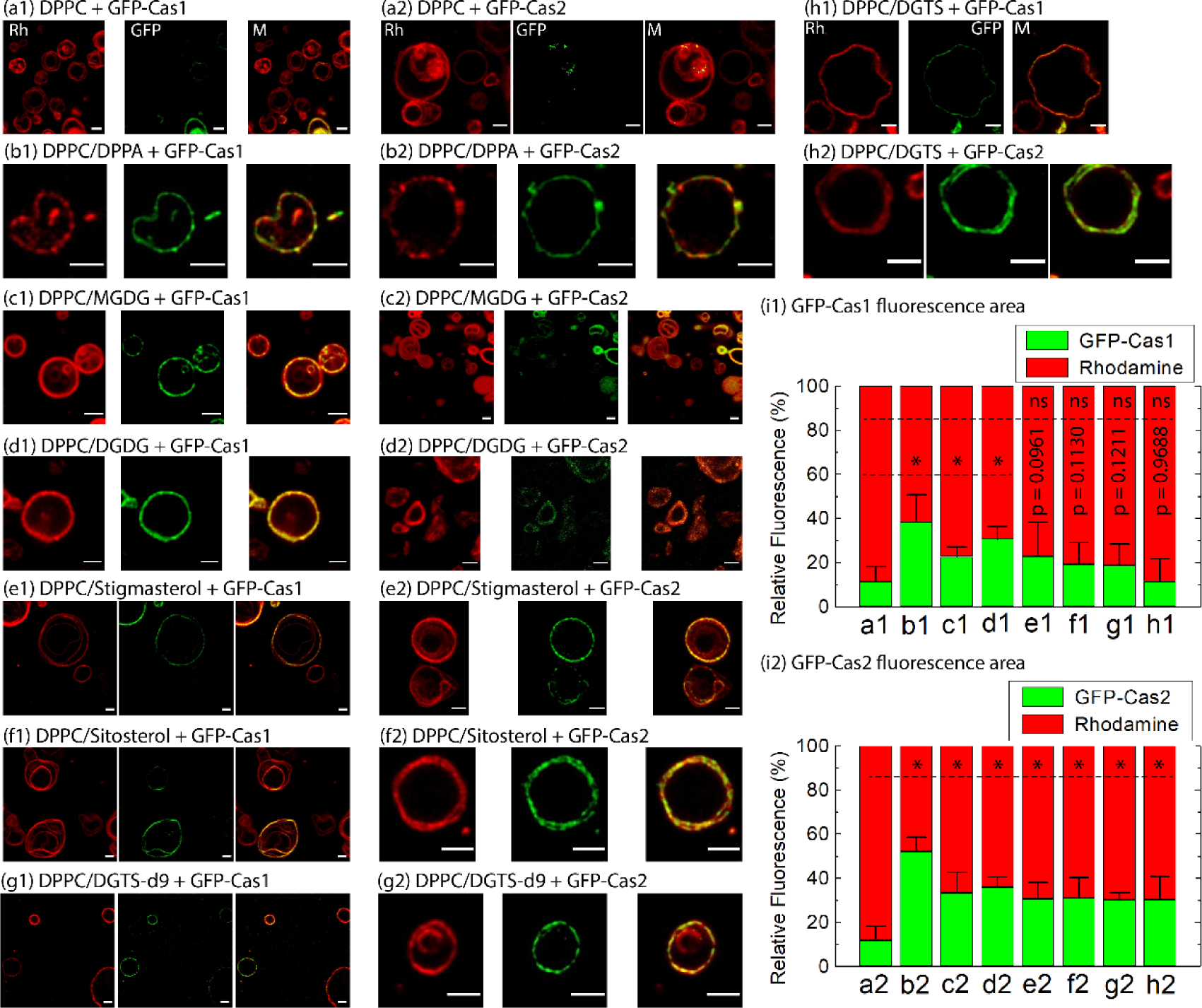
Confocal microscopic imaging to quantify the binding of GFP-Cas1 and GFP- Cas2 to GVs made of different lipids available in plants. The final concentration of GFP- Cas1 and GFP-Cas2 was 9.5 µM. From left to right (a1 – h1, a2 – h2), Rh denotes rhodamine-PE, GFP denotes GFP-Cas1 or GFP-Cas2, M: merged. (i1 and i2) Quantification of relative fluorescence areas of GFP-Cas1 and GFP-Cas2 on various GVs described from a-h. The values are mean ± SD. GFP fluorescence areas on DPPC GVs were used as the controls for comparison with those on other GVs. Asterisks (*) show a significant difference (p ≤ 0.05, two-independent populations *t-test*). *p*-values are shown when the difference is not significant (ns) at the indicated level (p ≤ 0.05). Number of values in each population: 6 – 11. Bars: 5 µm. **<u>Related to</u> Tables 1, 2**.

**Table 2.** Quantification of relative fluorescence areas of GFP-Cas1 and GFP-Cas2 on different composite GVs. The values are mean ± SD. The two-independent populations *t- test* was used to examine the significant differences, and *p*-values and number of values (N) are shown in parentheses. GFP fluorescence areas on DPPC GVs were used as the controls for comparison with those on other GVs. **<u>Related to</u> Figure 4**.

| GVs Samples | Relative fluorescence areas of<br>GFP-Cas1 (mean ± SD) %<br>(*p* value, N) | Relative fluorescence areas of<br>GFP-Cas2 (mean ± SD) %<br>(*p* value, N) |
| --- | --- | --- |
| a1,a2 (DPPC) | 11.2 ± 6.9<br>(1.0000, 8) | 11.8 ± 6.4<br>(1.0000, 7) |
| b1,b2 (PPPC/DPPA) | 38.1 ± 12.7<br>(0.0002, 8) | 52.2 ± 6.4<br>(0.0000001, 7) |
| c1,c2 (DPPC/MGDG) | 22.5 ± 4.6<br>(0.0048, 7) | 33.2 ± 9.5<br>(0.0006368, 7) |
| d1,d2 (DPPC/DGDG) | 30.7 ± 5.7<br>(0.00003, 9) | 35.9 ± 4.6<br>(0.0000004, 10) |
| e1,e2 (DPPC/stigmasterol) | 22.7 ± 15.7<br>(0.0961, 8) | 30.6 ± 7.5<br>(0.0002098, 9) |
| f1,f2 (DPPC/sitosterol) | 19.2 ± 9.9<br>(0.1130, 8) | 30.9 ± 9.3<br>(0.0004778, 10) |
| g1,g2 (DPPC/DGTS-d9) | 18.6 ± 9.7<br>(0.1211, 8) | 30.0 ± 3.4<br>(0.0001303, 6) |
| h1,h2 (DPPC/DGTS) | 11.4 ± 10.3<br>(0.9688, 7) | 30.3 ± 10.5<br>(0.0011800, 11) |

### Cytotoxicity of Cas1 and Cas2 on rubber leaves

To examine the cytotoxicity of Cas1 and Cas2 on rubber leaves at the cellular level, this study addressed the second question of whether Cas1 and Cas2 could directly damage and cause necrosis in rubber leaf tissues. Here, we tested the cytotoxicity of Cas1 and Cas2 on freshly detached young rubber leaves of highly susceptible clones (*H. brasiliensis* RRIV 1 and RRIV 4) and weakly susceptible clones (*H. brasiliensis* PB 255), as reported previously (Hieu et al., 2020; Nguyen et al., 2014). In these experiments, 6 – 10 small spots on a rubber leaf’s lower lamina side were slightly scraped on the cuticle surface before applying 0.083 µg Cas1 or Cas2 to each spot. We evaluated the cytotoxicity of Cas1 and Cas2 on rubber leaves by observing the color change and the size change in the lesion spots. As seen in **Figure 5**, neither Cas1 nor Cas2 caused any noticeable damage or color change in the infected spots of all rubber leaves after incubation for 24 h. However, a green to brown color change was seen in some leaf lesion spots of susceptible RRIV 1 and RRIV 4 rubber clones infected with Cas1 and Cas2 after incubation for 72 h. The necrotic areas in the leaf lesion spots were largely expanded and seen after incubation for 120 h (**Figures 5a1, a2, b1, b2**). In the case of the PB 255 rubber clone, necrosis in the leaf lesion spots was developed and seen after treatment with Cas1 for 72 and 120 h (**Figure 5c1**). However, necrosis in the leaf lesion spots developed very slowly and was not clearly seen after treatment with Cas2 for 120 h (**Figure 5c2**). This observation suggests that the cytotoxicity of Cas1 and Cas2 on the rubber leaves of a PB 255 clone was different. Noticeably, some leaf spots infected with Cas1 changed from green to brown and black colors (see black circles in **Figures 5a1, b1, c1**), while those infected with Cas2 only changed from green to brown color (**Figures 5a2, b2**). The blackened necrosis seen in lesion spots infected with Cas1 implies that Cas1 damaged leaf tissues more severely than Cas2. Once the damage of Cas1 and Cas2 on leaf tissues began, leaf tissue necrosis continuously developed and expanded in large areas. The cytotoxicity assay on rubber leaves (RRIV 4 clone) with fungus *C. cassiicola* showed that the necrotic leaf tissues surrounding filamentous fungi were produced and clearly seen at day5 and day7 post inoculation (see brown halos, **Figure 5d**). This result suggest that Cas toxins were made and secreted from fungus to damage the surrounding leaf tissues. Overall, we demonstrated that both Cas1 and Cas2 could directly cause necrosis in the leaves of some rubber clones. Our report also supports the previous finding of the cytotoxicity of Cas1 produced in *C. cassiicola* resulting in necrosis in rubber leaves (Ribeiro et al., 2019). Indeed, the cytotoxicity of Cas2 on rubber leaves has never been reported before. Here, we additionally report that Cas2 directly damaged and caused necrosis in rubber leaves.

**Figure 5.**
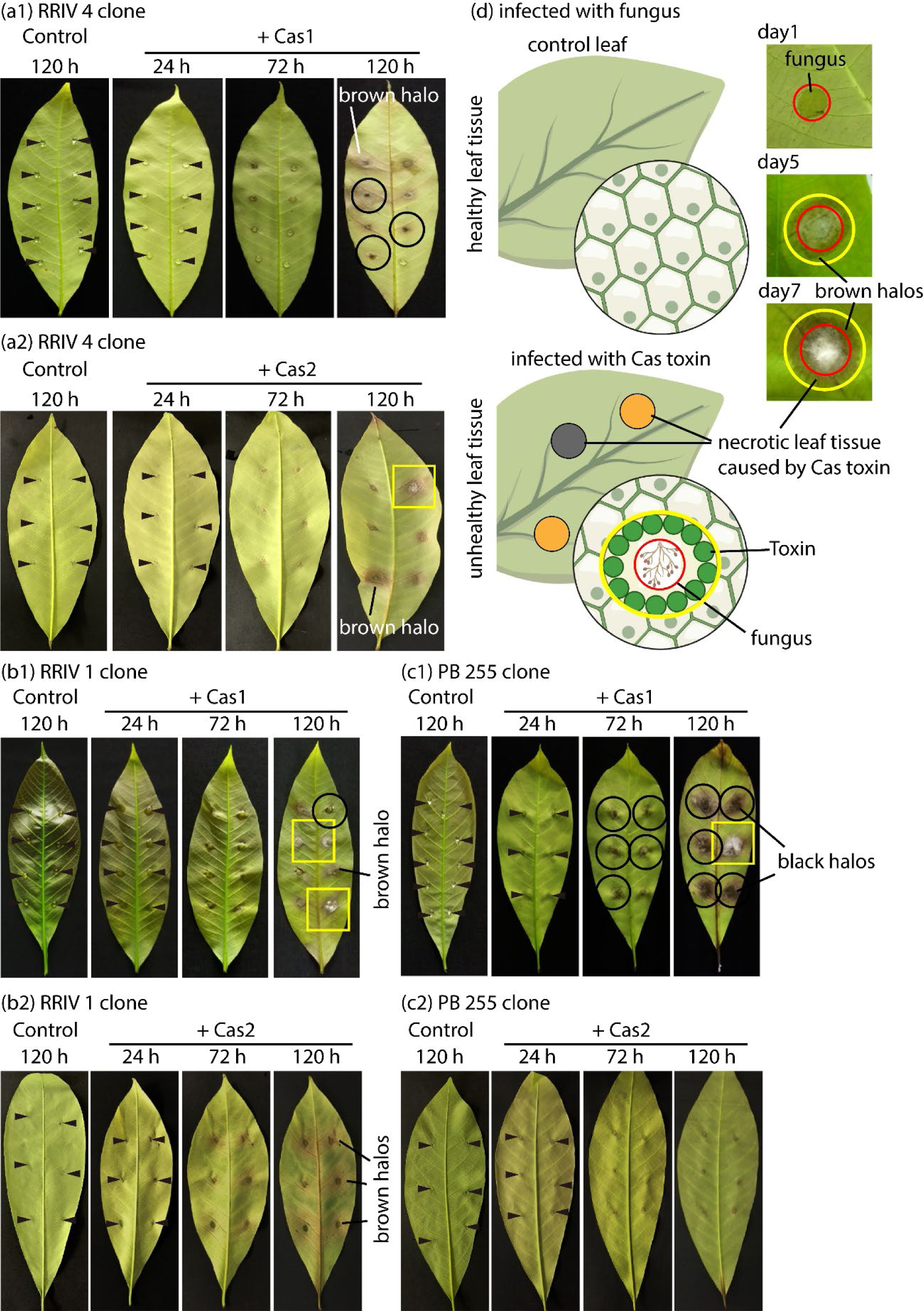
Cytotoxicity of rubber leaves of RRIV 4, RRIV 1, and PB 255 clones caused by Cas1 and Cas2. Six to ten small spots on the lower lamina sides of the healthy rubber leaves were slightly scraped before cytotoxicity tests. (a1 – c2): Control: Ten microliters of sterilized water were applied to each spot. Infected leaves: Ten microliters of 0.0083 µg/µl (∼0.083 µg) Cas1 or Cas2 were applied to each spot. Control and infected leaves were separately incubated under the same humidity (∼80 – 90%) and temperature (26 – 28°C) and photographed fter incubation at 24, 72, and 120 h by a conventional digital camera. (d) Rubber leaves (RRIV 4 clone) were infected with 10 µl conidial solution of fungus *C. cassiicola* (2x10^3^ spores/ml) and inoculated at 25°C and humidity (∼80 – 90%). Arrowheads: Initial positions of the infected spots. Black circles: blackened lesion spots: Rectangles: possible contamination of fungus. In (d), red circles: *C. cassiicola* growing areas; space between red and yellow circles: brown halos of necrotic leaf tissues caused by Cas toxin secreted from fungus. **<u>Related to</u>** Figure 6 and Figure 6-figure supplement 1

### Dissecting cellular structures of rubber leaves using cryo-SEM

This study addressed the third question of how Cas toxin could damage cellular and tissue structures in rubber leaves. To answer this question, we used low voltage cryo-SEM to anatomize the cellular and tissue structures in the leaf lesion spots developed in rubber leaves of susceptible RRIV 4 clones after infection with Cas1 for ∼120 h and compared them to the structures in healthy control leaves. Indeed, ultralow-temperature and low-voltage cryo-SEM has been proven to be a powerful technique for dissecting pristinely fractured biological cells without conductive metal coatings (Osumi et al., 2006).

The lamina, cuticle, upper and lower epidermis, palisade mesophyll, and spongy mesophyll in the healthy control leaves preserved their healthy cellular structures, which were identified easily (**Figures 6A, B, C, D, and Figure 6-figure supplement 1**). Additionally, intact cell walls and cytoplasmic membranes could be seen clearly in the fractured leaf tissues (**Figures 6C, D, and Figure 6-figure supplement 1**). In contrast, Cas1 strongly damaged the cellular and tissue structures in leaf lesion spots (**Figures 6E and Figure 6-figure supplement 1**). The upper and lower epidermis, spongy, and palisade parenchyma were devastated by Cas1. The cellular boundaries, such as cell walls and cytoplasmic membranes, in the fractured leaf tissues were hardly distinguished. Strikingly, large areas of cytoplasmic membranes in palisade mesophyll were found to be severely devastated by Cas1. The binding of Cas1 clusters, pore formation, biofilm-like formation, and expansion of Cas1 clusters at different degrees in the cytoplasmic membranes were clearly observed (see white rectangles in **Figure 6E, Figure 6-figure supplement 1**). Our results strongly suggest that cytoplasmic membranes were highly vulnerable to the pathogenicity of Cas1. This mechanism may also be similarly applied to understand the cytotoxicity of Cas2 on the leaves of susceptible rubber clones, although it needs to be verified.

**Figure 6.**
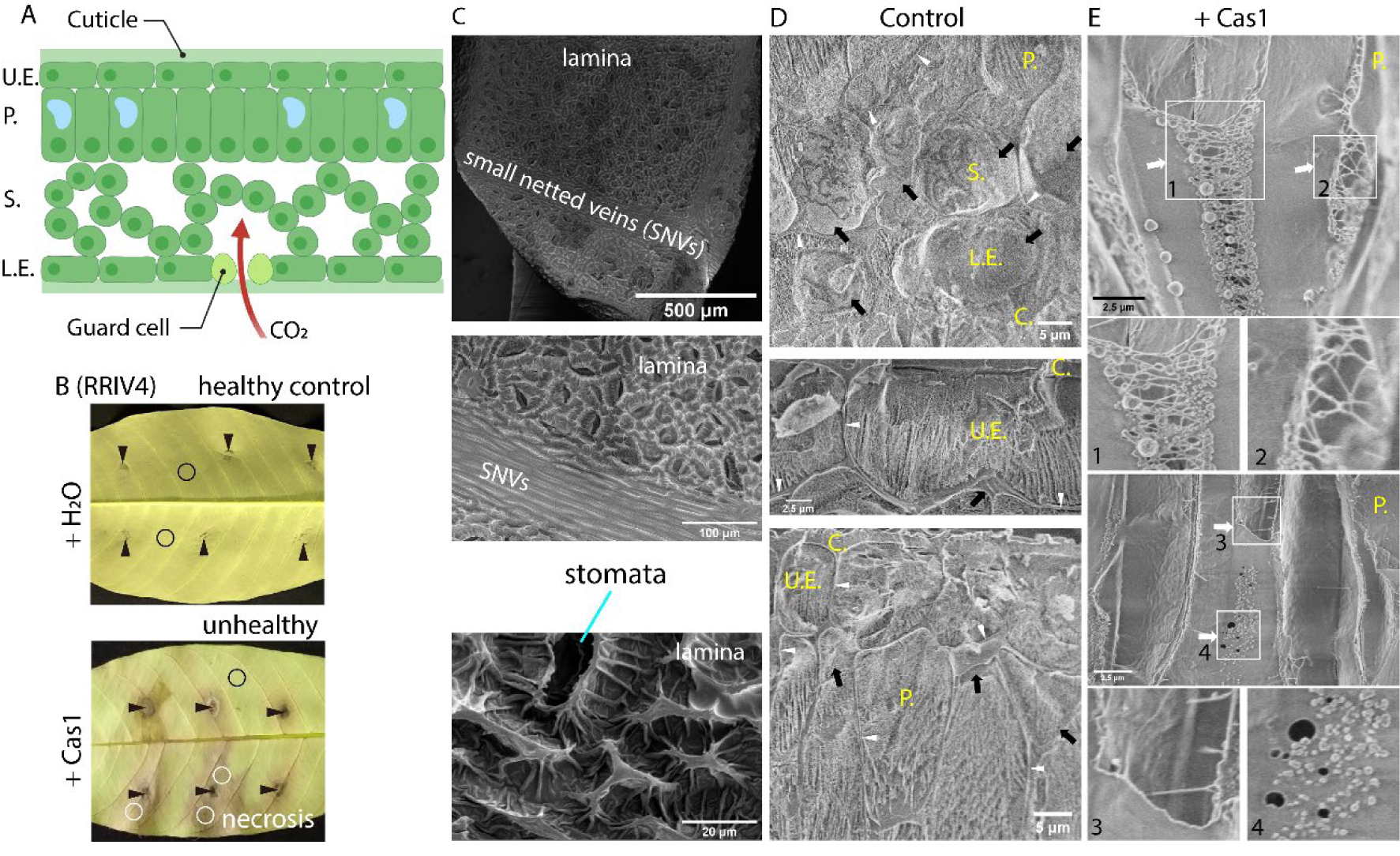
Cryo-SEM images of cellular and tissue structures of healthy controls and lesion spots in rubber leaves (RRIV 4 clone) caused by Cas1. (**A**) An illustration of leaf tissue anatomy; (**B**) Lower lamina side of the healthy control and infected rubber leaves (+ 0.083 µg Cas1) after incubation for ∼ 120 h. The small areas of the healthy control (black circles) and lesion spots (white circles) in rubber leaves shown in B were punched out and plunged into liquid ethane, and cellular and tissue structures were dissected. Black arrowheads: initially infected spots. (**C)** Lower lamina side of rubber leaves. (**D**, **E**) Cellular and tissue structures of the healthy control and lesion spots, respectively. Black and white arrows denote the healthy and damaged cytoplasmic membranes, respectively. White arrowheads: the healthy cell wall. C: cuticle, U.E.: upper epidermis, L.E.: lower epidermis, P.: palisade mesophyll, S.: spongy mesophyll. White rectangles denote biofilm-like structures of Cas1 formed on cytoplasmic membrane. **<u>Related to</u>** Figure 5 and Figure 6-figure supplement 1

## Discussion

### Cytoplasmic membrane vulnerability to Cas toxin plays a crucial role in HST interactions

Cytoplasmic membranes are essential in all living cells. In plant cells, life strongly relies on the presence of a healthy functional cytoplasmic membrane enveloping the cytosol, thereby forming a boundary between the internal and external cytosolic environments. The cytoplasmic membrane plays pivotal roles in protecting cytosolic contents, such as the chloroplast, Golgi apparatus, mitochondria, endoplasmic reticulum, and nucleus, and furnishing a sophisticated lipid environment for various vital activities across membranes, such as the transport activities of protein channels and transporters (Assmann and Haubrick, 1996; Kang et al., 2011; Maure et al., 2008). Phospholipids (PC, PA), glycerolipids (MGDG, DGDG), sterols (stigmasterol, sitosterol), and betaine lipids (DGTS-d9, DGTS) are present in the cytoplasmic membranes of plant cells (Bae et al., 2020; Dörmann and Benning, 2002; Dufourc, 2008a, 2008b; Hartmann, 1998; Liengprayoon et al., 2011; Mackender and Leech, 1974; Murakami et al., 2018; Poincelot, 1976)^,53^. The vulnerability of cytoplasmic membranes to Cas1 has been reported previously (Barthe et al., 2007; de Lamotte et al., 2007). When observing the ultrastructures of the necrotic leaf tissues caused by Cas1, they found that Cas1 severely damaged the plasma membrane. Plasmolysis (e.g., membrane permeability) was modified. The neutral deuterated (^2^H)DPC membrane was weakly susceptible to Cas1, although they could not rule out the interaction of Cas1 with other lipid classes (Barthe et al., 2007). In this study, we showed that both Cas1 and Cas2 could directly damage and cause necrosis in the leaves of some susceptible rubber clones. Our electron microscopic analyses strongly suggest that cytoplasmic membranes were highly vulnerable to the pathogenicity of Cas toxin (**Figures 6E and Figure 6-figure supplement 1**), supporting the previous finding that Cas1 could damage cytoplasmic membranes (Barthe et al., 2007; de Lamotte et al., 2007). We also explored here that the negative phospholipids (DPPA), glycerolipids (MGDG, DGDG), and sterols (stigmasterol, sitosterol) available in cytoplasmic membranes play key roles in the binding and disruption activity of Cas1 and Cas2 to the cytoplasmic membrane. Neutral phospholipids (DPPC) and betaine lipids (DGTS-d9, DGTS) are indeed weakly sensitive to the disruption activities of Cas1 and Cas2. Our finding is crucial to understand the binding and disruption selectivity of Cas toxin to different lipid compositions in cytoplasmic membranes.

Studying the HST interaction of Cas isoforms with lipid membranes is crucial to understand the host-selective pathogen in CLF disease. Currently, the exact mechanism remains unclear. In general, mycotoxins are categorized as host-specific or nonspecific toxins. Some plant pathogenic fungi can synthesize mycotoxins as secondary metabolites that act as virulence factors to develop severe fungal infections (Meena and Samal, 2019; Petrov et al., 2018; Takaoka et al., 2014). Several important organelles inside plant cells, such as the cytoplasmic membrane, chloroplasts, mitochondria, and endoplasmic reticulum, have previously been suggested as target sites of selective AK toxins (Tsuge et al., 2013). In particular, selective AK and fumonisin toxins were reported to damage or modify cytoplasmic membranes in various ways (Park et al., 1976; Riley et al., 1996). Here, we demonstrated that Cas1 and Cas2 directly damaged and caused necrosis in the leaves of susceptible rubber clones (RRIV 1, RRIV 4, and PB 255). RRIV 1 and RRIV 4 clones were sensitive to both Cas1 and Cas2 (**Figures 5a1, a2, b1, b2**). Strikingly, a PB 255 clone was strongly resistant to Cas2 but susceptible to Cas1 (**Figures 5c1, c2**). These results suggest that the cytotoxicity of the Cas1 and Cas2 isoforms varied in different rubber clones, depending on the susceptibility of cytoplasmic membranes to the binding and disruption of Cas toxin.

### Biofilm-like formation of Cas toxin and its physiological implication

Strikingly, we also disclose that Cas toxin can form biofilm-like structures onto specific lipid membranes composed of negative phospholipid (DPPA), glycerolipids (MGDG, DGDG), and sterol (sitosterol) (**Figures 1b1, b2, c1, d1, d2, and 3A, B, and Videos 3, 4, 5, 7, 8, 17, 18**). However, these biofilm-like structures are not formed on lipid membranes composed of betaine lipids (DGTS-d9) (**Figure 3C, Video 19**). Our realtime HS-AFM observation supports that the highly susceptible lipid membranes containing negative phospholipid, glycerolipids, and sterol can recruit the binding of Cas1 and Cas2 onto their surfaces to form biofilm-like structures. The biofilms are either covered on the membrane surface or partially incorporated onto the lipid membrane and cause severe damage on membranes. Most likely, the biofilm-like formation of Cas toxin also plays an important role in damaging cytoplasmic membranes, as seen in Cryo-SEM images (**Figure 6E, Figure 6-figure supplement 1**).

Finally, we hypothesized the molecular mechanism of the host-selective toxic interaction of Cas toxin with specific lipid membranes and its cytotoxicity on the susceptible rubber leaves (**Figure 7**). The brown and black necrotic halos of leaf tissues surrounding necrotrophic fungus *C. cassiicola* infected in rubber leaves are caused by the selective Cas toxin secreted during the growth and invasion of fungus. The Cas toxin binds to and strongly disrupts the specific and susceptible membrane lipids such as negative phospholipids, glycerolipids, and sterols available in cytoplasmic membranes of the specific plant cells. PA, MGDG, DGDG, and sitosterol lipids available in cytoplasmic membranes may play an important role to recruit the binding and formation of biofilm-like structures of Cas toxin onto cytoplasmic membranes. The selective binding and disruption activities of Cas toxin on specific cytoplasmic membranes cause plasmolysis and eventually cell death.

**Figure 7.**
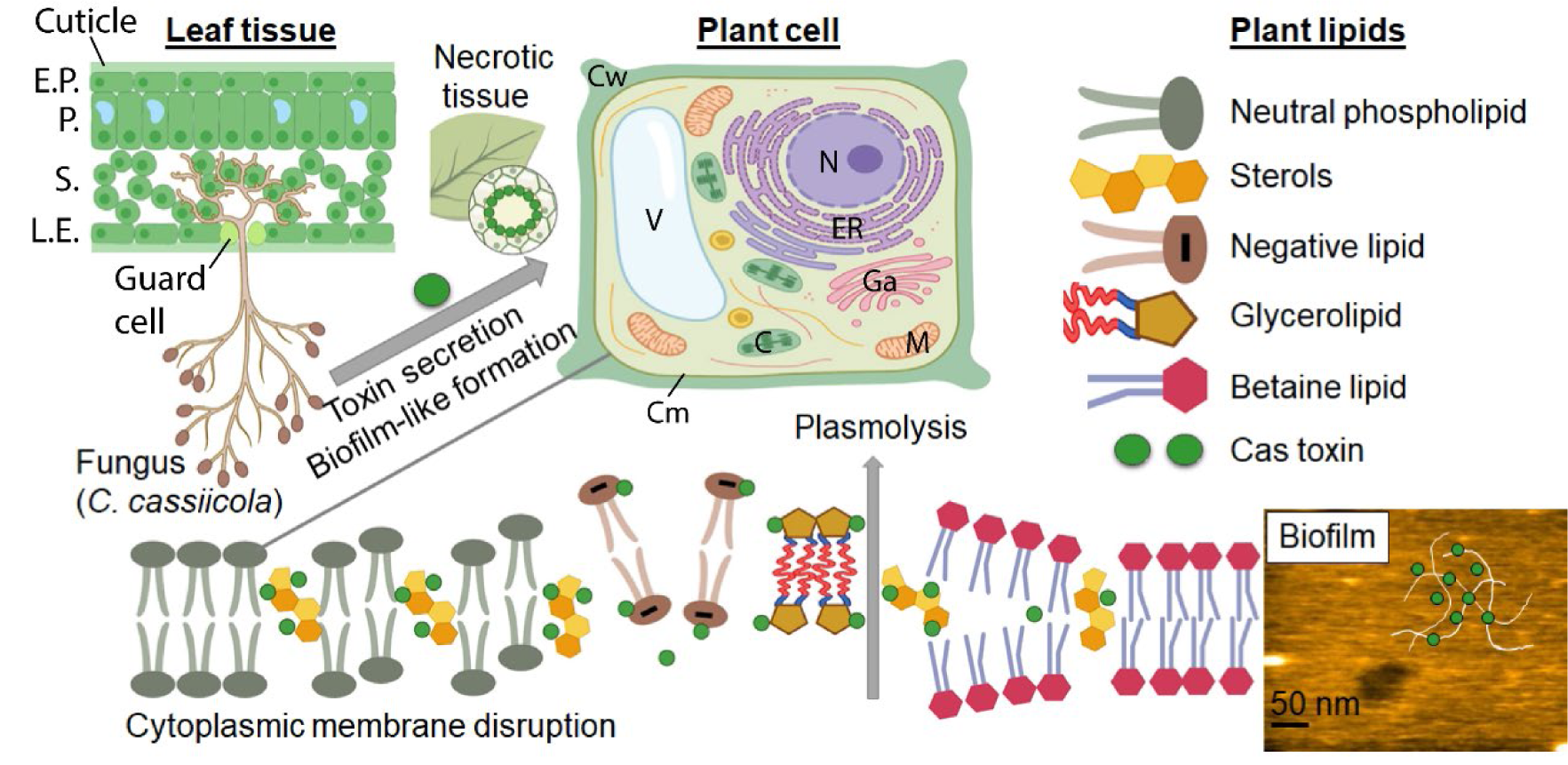
Hypothesis of the host-selective toxic interaction of Cas toxin with specific lipid membranes and its cytotoxicity arise from the formation of biofilm-like structures and disruption activities onto specific cytoplasmic membranes composed of susceptible plant lipids (i.e., negative phospholipids (PA), glycerolipids (MGDG, DGDG), and sterols (stigmasterol, sitosterol)). Cas toxin (Cas1, Cas2), Cm: cytoplasmic membrane, Cw: cell wall, N: nucleus, ER: endoplasmic reticulum, Ga: Golgi apparatus, C: chloroplast, M: mitochondria, V: vacuole, U.E.: upper epidermis, L.E.: lower epidermis, P.: palisade mesophyll, S.: spongy mesophyll.

## Methods

### Materials

DNA oligonucleotides encoding cassiicolin (Cas) genes (Cas1 and Cas2) of *Corynespora cassiicola* causing CLF disease in rubber trees (Déon et al., 2014, 2012b, 2012a) were purchased from IDT (Tokyo Japan). The PcoldGFP plasmid was a generous gift from Prof. Taro Q.P. Uyeda (Waseda University, Japan). All necessary primers used for PCR and cloning Cas1, Cas2, GFP-Cas1, and GFP-Cas2 were purchased from Macrogen Japan (Tokyo, Japan). The oligonucleotides and amino acid sequences of Cas1 and Cas2 used in this study are summarized in **Table S5**. All synthetic lipids available in plants were purchased from Avanti Polar Lipids (Alabama, USA). The structures and physical properties of these lipids are clarified and summarized in **Figure S3**.

### Protein preparation

The genes encoding Cas1 and Cas2 proteins were similarly inserted into the pColdI vector at the *KpnI* and *XbaI* sites (Ngo et al., 2015). Similarly, the PcoldGFP vector (generated two cloning sites by *BamHI* and *HindIII*) was employed to construct GFP- Cas1 and GFP-Cas2 proteins. The successfully constructed plasmids for the expression of Cas1, Cas2, GFP-Cas1, and GFP-Cas2 genes were confirmed by DNA oligonucleotide sequences. All recombinant proteins containing a 6xHis-tag on the N-terminus (**Table S5**) were designed for purification using a Ni-NTA resin column. For simplicity, we consistently designated these recombinant proteins as Cas1, Cas2, GFP-Cas1, and GFP- Cas2.

Cas1, Cas2, GFP-Cas1, and GFP-Cas2 were expressed in BL21 (DE3) *E. coli* according to the manufacturer’s instructions for PcoldI (Takara). Briefly, the transfected *E. coli* cells were grown in LB broth containing 100 µg/ml ampicillin and 10 µg/ml methyl 3-O(alpha-D-mannopyranosyl)-alpha-D-mannopyranoside (MDMDM) to an OD600 of ∼ 0.6. Then, the cell suspension was cooled to 15°C on ice. Protein expression was induced by adding 1 mM IPTG. The cell suspension was continuously fermented in a shaking incubator at 140 rpm and 15°C for 24 h. Then, the cells were harvested by low centrifugation (4000 rpm, 10 min, 4°C). The cell pellet was gently washed twice with cold 10 mM Tris-HCl buffer (pH 7.4). Next, the cell pellet was resuspended in lysis buffer containing 50 mM Tris-HCl buffer (pH 8.0), 300 mM KCl, 5 mM EGTA, 20 mM imidazole (pH 7.4), 1% (w/v) lauryl maltose neopentyl glycol (LMNG) (Anatrace), 10 mM β-mercaptoethanol, proteinase inhibitor cocktail (Sigma), and 1 mM PMSF. Cell lysis was carried out overnight in a rotary incubator at 4°C. Next, the cells were finally disrupted using a French press (repeated twice, ∼140 MPa). After low centrifugation (15,000 rpm, 15 min, 4°C), the crude protein solution was finally collected by ultracentrifugation (40,000 rpm, 30 min, 4°C) and subsequently incubated with Ni-NTA (GE healthcare) resin for 2 h at 4°C. The resin was recovered and transferred into a clean plastic column for washing. The resin was washed carefully in cold wash buffer containing 10 mM Tris-HCl (pH 7.4), 20 mM imidazole (pH 7.4), 1 M NaCl and 10 mM β-mercaptoethanol until proteins in the washout solution were undetectable. Elution of proteins was carried out with elution buffer containing 10 mM Tris-HCl (pH 7.4), 800 mM imidazole (pH 7.4) and 10 mM β-mercaptoethanol. The eluate was dialyzed against dialysis buffer containing 10 mM Tris-HCl (pH 7.4), 50 mM KCl, 50% glycerol, and 10 mM β-mercaptoethanol using a dialysis membrane (MWCO 3300 Da) overnight at 4°C. The dialysate was recovered and concentrated and finally subjected to low centrifugation (15,000 rpm, 10 min, 4°C) to remove any protein aggregates. Protein concentration was measured by the Bradford method or UV absorption (Cas1: ε280 = 8855 M^-1^. cm^-1^ or A280 = 0.984 (1 mg/ml), 9003.27 Da; Cas2: ε280 = 8855 M^-1^. cm^-1^ or A280 = 0.981 (1 mg/ml), 9025.28 Da; GFP-Cas1: ε280 = 29380 M^-1^. cm^-1^ or A280 = 0.830 (1 mg/ml), 35400.95 KDa; GFP-Cas2: ε280 = 29380 M^-1^. cm^-1^ or A280 = 0.815 (1 mg/ml), 35422.07 KDa). Small aliquots of proteins were snap-frozen in liquid nitrogen and stored at -80°C. The activities of target proteins were characterized by common biochemical assays, mass spectrometry, HS-AFM, confocal microscopy, cryo-SEM, and cytotoxicity assays on rubber leaves.

### Giant liposome vesicles

We made giant liposome vesicles (GVs) from different lipids available in plants for imaging the binding of GFP-Cas1 and GFP-Cas2 to different lipid membranes using confocal microscopy. The electroformation methods for making GVs have been well established in several previous studies (Angelova and Dimitrov, 2007, 1986; Dimitrov and Angelova, 1988). The lipid compositions are described in **Table S3** and clarified in **Figure S3**. Briefly, the lipid mixtures dissolved in chloroform were stored at -20°C. The mixtures were warmed at room temperature for 10 min. A 20 µl lipid mixture was thoroughly spread on indium tin oxide (ITO)-coated glass slides (Nanion, Germany). The dried thin film was formed by the complete evaporation of chloroform at room temperature for approximately 10 min. To make GVs on the ITO-coated glass slide, we used a Vesicle Prep Pro (Nanion, Germany). An O-ring with a diameter of 16 mm was used to make a chamber furnishing the dried thin film inside. A 250 µl filter sterile electroformation buffer containing 1 M D-sorbitol, 1 mM EGTA, and 2 mM HEPES-NaOH (pH 7.4) was applied into a chamber sandwiched between two conductive sides of the ITO-coated glass slides. The protocol for making GVs was set up as follows: Initial (0 V, 5 Hz, 50°C, 5 min); Start (3 V, 10 Hz, 50°C, 10 min); Main (3 V, 10 Hz, 50°C, 60 min); Fall (0 V, 10 Hz, 50°C, 5 min); End (0 V, 10 Hz, 20°C). The formed GVs were gently collected and stably stored at 4°C for approximately 1 – 2 weeks.

### Confocal microscopy

The FluoroDish cell culture dishes (FD35-100, WPI) were cleaned by dipping in 3 M KOH solution for 1 h, subsequently washed in Milli-Q water approximately 10 times, dried at 60°C for 1 h or more, and stored at room temperature before use. GV solution (∼0.8 mM lipid) was diluted in imaging buffer containing 5 mM HEPES-NaOH (pH 7.4), 5 mM NaCl, 0.05 mM EDTA, and 1 M D-sorbitol to a final lipid concentration of ∼0.2 mM. To assay the binding of Cas1 or Cas2 to different lipid membranes, we gently mixed 9.5 µM GFP-Cas1 or GFP-Cas2 with 100 µl of diluted GV solution, incubated the mixture at room temperature for 5 min, and subsequently introduced the mixture into a cleaned FluoroDish. In most cases, we could image the GVs weakly absorbing onto the glass surface by gravity force. Only in the case of negatively charged DPPC/DPPA GVs did we further added 300 mM NaCl and 10 mM MgCl2 into the imaging buffer for better absorption of these GVs onto the glass surface.

Confocal imaging experiments were performed in a STED confocal microscopy (Abberior Instruments, Germany). We used a 100x oil-immersed objective lens (NA=1.46). Images were taken using Imspector software. Two channel images were acquired one by one for each pixel at 50 x 50 x50 nm resolution per xyz pixel. The confocal imaging conditions were set up as follows: SiR channel (Ex: 652 nm) for rhodamine, EGFP channel (Ex: 489 nm) for GFP, 15% laser power source used for both channels. The images were analyzed using Fiji-ImageJ (NIH, USA). Initially, we corrected the background for both channels. After a Gaussian-blur filter (sigma radius = 2.0) was applied, the images were converted to binary images with a minimum threshold set value of 12 for measuring the pixel areas in each channel. The rhodamine area where Cas is not bound was calculated by subtracting the GFP channel from the rhodamine channel. Then, the relative fluorescence area was calculated for a quantitative comparison of the binding area of GFP-Cas1 and GFP-Cas2 to different GVs.

### Mass spectrometry

The target Cas1 protein band was excised from the electrophoresis gel and subjected to in-gel digestion (Hellman et al., 1995). The peptide fractions were analyzed by a solariX (Bruker Daltonics, Billerica, USA), as previously described (Kawakita et al., 2016).

### High-speed atomic force microscopy

We used a laboratory-built high-speed atomic force microscope (HS-AFM) as described previously (Ando et al., 2008; Ngo et al., 2015; Uchihashi et al., 2012). HS-AFM imaging was carried out in solution using tapping mode with small cantilevers (BL- AC10DS-A2, Olympus, Tokyo, Japan) whose spring constant, resonant frequency in water, and quality factor in water were ∼0.1 N/m, ∼500 kHz, and ∼1.5, respectively. Our lab-made ferrocene tip on an original tip-less cantilever was grown through electron beam deposition (EBD) using scanning electron microscopy (ZEISS Supra 40 VP/Gemini column, Zeiss, Germany). Typically, the EBD tip was grown under vacuum (1 – 5 x 10^-6^ Torr), aperture size (10 µm), and EHT voltage (20 kV) for 30 s. The EBD ferrocene tip was further sharpened using a radio frequency plasma etcher (Tergeo Plasma Cleaner, Pie Scientific, CA, USA) under an argon gas atmosphere (typically at 180 mTorr and 20 W for 30 s). During HS-AFM imaging, the free-oscillation peak-to-peak amplitude of the cantilever (*A0*) was set to ∼1.6 – 1.8 nm, and the feedback amplitude set-point was set at ∼0.9*A0*.

### Real-time imaging of disruption of lipid membranes caused by Cas1 and Cas2

We prepared small unilamellar vesicles (SUVs) or liposomes by the thin-film hydration method and mica-supported lipid bilayer as described previously (Ngo et al., 2015). In this study, DPPC, DPPC/DPPA (80/20 wt%), DPPC/MGDG (80/20 wt%), DPPC/DGDG (60/40 wt%), DPPC/stigmasterol (80/20 wt%), DPPC/sitosterol (80/20 wt%), DPPC/DGTS-d9 (80/20 wt%), and DPPC/DGTS (80/20 wt%) liposomes were made. Typically, liposomes were made in Milli-Q water at 1 mg/ml and stored at −20°C. To form a lipid bilayer on mica, liposomes were diluted in 10 mM MgCl2 to 0.1 mg/ml and sonicated with a bath sonicator at room temperature (AUC-06 L, AS ONE, Osaka, Japan) for 1 min. Then, 2 µl of the sonicated liposomes was deposited on the surface of freshly cleaved mica (2 mm in diameter), which had been glued onto a glass stage beforehand, and incubated for approximately 2 h at room temperature (24 – 26°C) in a humid sealed container to avoid surface drying.

Real-time imaging of the binding and disruption activities of Cas1 and Cas2 to lipid membranes formed on mica was carried out in C-buffer containing 10 mM HEPES-NaOH (pH 7.4), 0.1 mM EDTA, and 100 mM NaCl at room temperature by using HS-AFM. In all HS-AFM experiments excepted noted, we consistently used 0.95 µM Cas1 or Cas2 as a final concentration to examine the susceptibility of different lipid membrane compositions to Cas1 and Cas2. The sample volume in the HS-AFM imaging experiment was ∼70 µl.

### Cytotoxicity on rubber leaves caused by Cas1 and Cas2

Young and healthy leaves of rubber trees (*H. brasiliensis* RRIV 4, RRIV 1, and PB 255 clones) grown in southern Vietnam were freshly collected in the early morning. It usually took 4 – 5 h before a cytotoxicity test. The leaves were carefully disinfected with 60% ethanol and sterilized Milli-Q water. Three rubber leaves were placed on a plastic net under one experimental condition and separately incubated inside a plastic box (40 cm x 40 cm x 5 cm). The moisture was maintained by laying down wet Whatman filter paper under a plastic net, avoiding direct contact with leaves. To test the cytotoxicity of Cas1 and Cas2 on rubber leaves, 6 to 10 small spots on the lower lamina sides of leaves were chosen and slightly scraped on the cuticle wax layer with a small pipette tip. Next, 10 µl of 0.0083 µg/µl (∼ 0.083 µg) Cas1 or Cas2 was applied to each spot. The molar concentration of Cas1 and Cas2 infected in each spot was kept almost the same as that used for HS-AFM observation, although the final sample volume used in HS-AFM observation was ∼ 7 times higher. The control experiments were performed similarly by applying 10 µl of sterilized Milli-Q water to each spot. The control and infected leaf samples were separately incubated and maintained under the same humidity (∼80 – 90%) and temperature (∼26 – 28°C) in the dark. For sample analyses, leaves in each moist plastic box were carefully photographed after 24, 72, and 120 h of infection by using a conventional digital camera. Each experimental condition was repeated 2 – 3 times. The leaf lesion spots caused by Cas1 and Cas2 on different rubber clones were compared by observing color changes and lesion size and dissecting the cellular structures by cryo-SEM.

Pathogenicity test on rubber leaves with *C. cassiicola*: Healthy rubber leaves (RRIV 4 clone) were washed with 0.2% sodium benzoate for 30 seconds, then followed by rinsing three times with sterile distilled water. After draining the leaves with sterile paper, rubber leaves were put up side down on the wire mesh in plastic containers in order to separate the leaves and moisture absorbent paper. The infection on rubber detached leaves was conducted by dropping 10 μl of *Corynespora cassiicola* conidial solution with the concentration of 2x10^3^ spores/ml at 8 points per leaf using micro pipette. Next, the plastic containers were placed in a room at 25°C and humidity (80 – 90%) under 12 hours of fluorescent light per day within six days. The pathogenicity of *C. cassiicola* was evaluated and recorded based on the modified method by Nghia et al., 2008.

### Cryo-SEM imaging

We selected the healthy (control, + water, 120 h) and infected (+ Cas1, 120 h) leaves of one rubber clone (RRIV 4) to anatomize the leaf tissues and cellular structures. To make the Cryo-SEM samples, healthy and lesion-containing areas surrounding the initially scraped spots were chosen. The ultrastructures of leaf tissues were analyzed by low voltage cryo-scanning electron microscopy (Cryo-SEM), a powerful technique for imaging pristinely fractured biological cells without conductive metal coating (Osumi et al., 2006). Leaf samples were punched out with a 3 mm diameter circular blade and plunged into liquid ethane at 90 K. Cryo-SEM images were obtained from a Helios G4 CX (Thermo Fisher Scientific, USA) with a PP3010 Cryo-SEM transfer system (Quorum Technologies, UK). Frozen leaf samples were clamped perpendicular at the top and the center of a metal cylinder on the transfer shuttle and fractured inside the vacuum chamber to face the freshly fractured depth profile just on the top of the cylinder without conductive metal coating. No ice contamination growth was visible during the fracture of samples under high vacuum conditions below a pressure of 5.0 x 10^-5^ Pa. The sample stage temperature was kept at 93 K. A 1 kV electron beam of the landing energy with a stage bias voltage of 2 kV was used substantially in low voltage cryo-SEM imaging of the fractured sample surfaces embedded in a cylindrical shape holder. Notably, only when the lower lamina side of rubber leaves was imaged we coated the leaf surface with platinum in the vacuum chamber before imaging.

### Analyses of HS-AFM images

HS-AFM images were viewed, analyzed, and exported by using the laboratory built software Kodec4.4.7.39. Kodec4.4.7.39 software was coded in Visual C# (Visual Studio 2010, Microsoft, USA), and source code 1 is openly available (Ngo et al., 2015). All filters and subroutines for image analyses used in the present study were included in the software. We confirmed the compatibility between the software and computers operated with Windows 8 or 10. This software should be cited as Sakashita M, M Imai, N Kodera, D Maruyama, H Watanabe, Y Moriguchi, and T Ando. 2013. Kodec4.4.7.39. Briefly, AFM images were flattened and filtered (Gaussian blur, Sigma radius: 1 nm) to increase the S/N ratio. AFM images were converted and extracted to BMP format images. The image sequences were converted to a Video format (AVI) using Fiji-ImageJ software (NIH, USA).

### Artworks

All illustrative graphics used in this study were done by using Biorender, Adobe Illustrator 2021 (Adobe, USA). Data figures were made by Origin Pro 2016 graphic software (OriginLab, USA) and Igor Pro graphic software (Wavematrics Inc., USA).

### Statistics

All data (mean ± SD) were statistically processed using an Origin Pro 2016 graphic software (OriginLab, USA). The differences were statistically tested using two-independent populations *t-test*.

## Acknowledgements

This work was partly supported by FY2018 and FY2019 Bio-SPMs Collaborative Research of WPI Nano Life Science Institute (WPI-NanoLSI), Kanazawa University, Japan to NBQ and KXN, and partly funded from Ho Chi Minh City Open University (No. E2019.05.1), Vietnam to NNBC; Grants-in-aid from the Ministry of Education, Culture, Sports, Science and Technology, Japan to KXN (No. 19K06581). We deeply thank Prof. Taro Q.P. Uyeda (Waseda University) for his generous gift of a plasmid. We thank Dr. Thien Quang Huynh (Kanazawa University, Vietnam Academy of Science and Technology) for sample preparation and data analysis.

## Author contributions

KXN and NBQ designed concepts. KXN designed and constructed plasmids, and KXN, PDNN, TOM expressed, purified, and analyzed proteins. NBQ, PDNN, NNBC, NPV, and NAN performed the cytotoxicity test. KXN and PDNN performed HS-AFM imaging. HF, KXN, and NBQ performed cryo-SEM observation and analyzed data. TS, MM, and KXN performed mass spectrometry and analyzed data. KXN, TI, and TF performed confocal microscopy. KXN, PDNN, NK, TA, HK, and NBQ analyzed HS-AFM data. KXN, NBQ, and TA wrote the manuscript.

## Nomenclature

DPPC: 1,2-dipalmitoyl-sn-glycero-3-phosphocholine DPPA: 1,2-dipalmitoyl-sn-glycero-3-phosphate MGDG: Monogalactosyldiacylglycerol
DGDG: Digalactosyldiacylglycerol
DGTS-d9: 1,2-dipalmitoyl-sn-glycero-3-O-4’-[N,N,N-trimethyl(d9)]-betaine DGTS: 1,2-dipalmitoyl-sn-glycero-3-O-4’-(N,N,N-trimethyl)-betaine
GVs: giant liposome vesicles
Cas: cassiicolin
CLF: corynespora leaf fall
CCP: pathogenic *C. cassiicola* isolates in rubber trees originally from the Philippines HS-AFM: High-speed atomic force microscopy
Cryo-SEM: cryo-scanning electron microscopy
MALDI FT-ICR: matrix-assisted laser desorption/ionization Fourier-transform ion cyclotron resonance
IPTG: Isopropyl β-D-1-thiogalactopyranoside PMSF: Phenylmethylsulfonyl fluoride MWCO: Molecular weight cutoff
RRIV: Rubber research institute of Vietnam
PB: Prang Besar (a name of a rubber research center in Malaysia)

## Figure supplements

**Figure 1-figure supplement 1.**
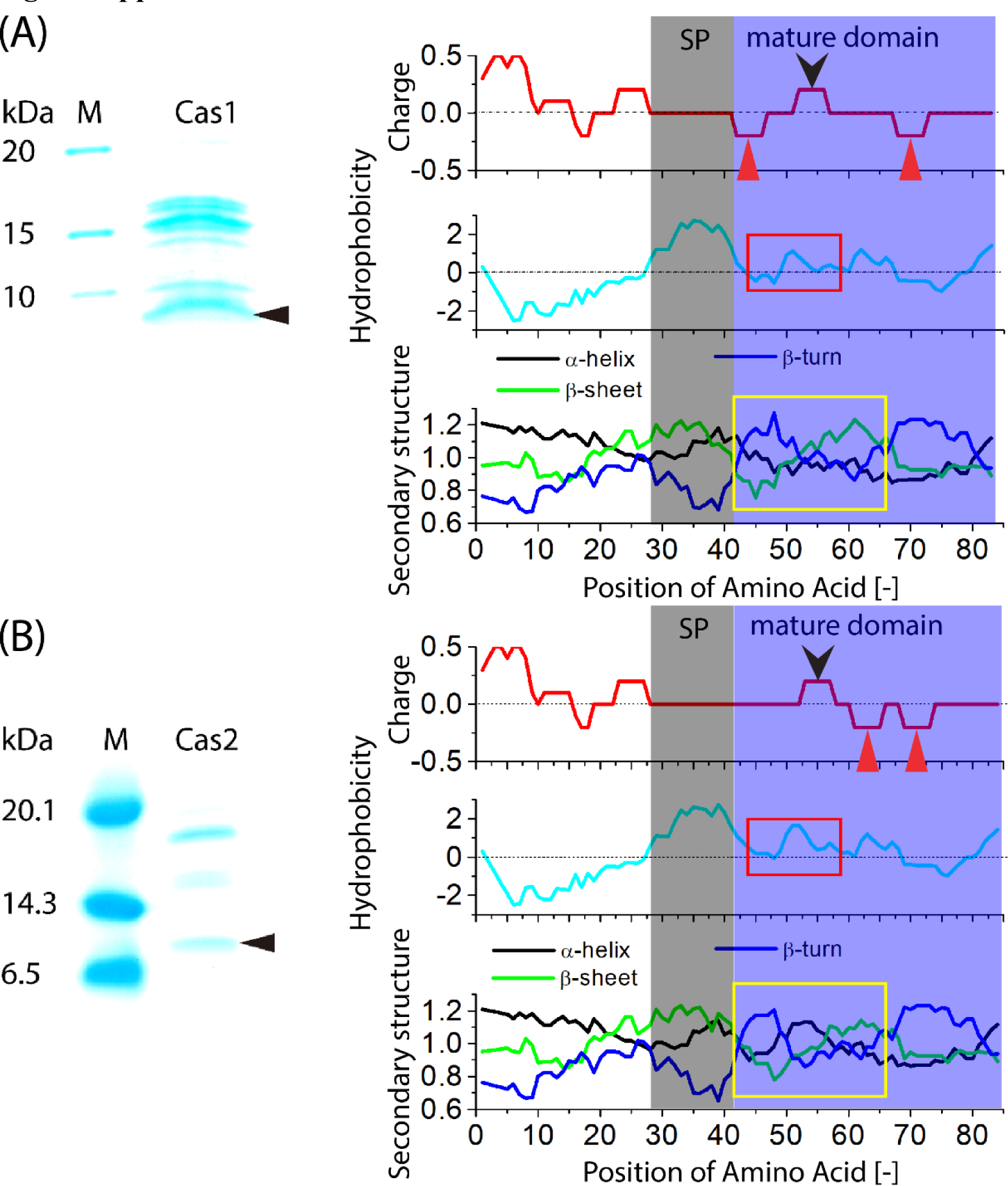
Purification of Cas1 and Cas2 from *E. coli*. (**A**) SDS-PAGE gel images of Cas1 and Cas2 stained with Coomassie Brilliant Blue R-250 (CBB) (**B**) Computing charge (http://www.bioinformatics.nl/cgi-bin/emboss/charge), hydrophobicity, and secondary structure (alpha-helix, beta-sheet, and beta-turn) scales based on the profiles produced by amino acid sequences of Cas1 and Cas2 (https://web.expasy.org/protscale/). SP: signal peptide; red and black arrowheads show the positions of the negatively and positively charged amino acids, respectively, in mature domains; red and yellow rectangles show the regions in which the hydrophobicity and secondary structure of Cas1 and Cas2 may differ. **<u>Related to</u> Figure 1 and Figure 1-table 1.**

**Figure 1-figure supplement 2.**
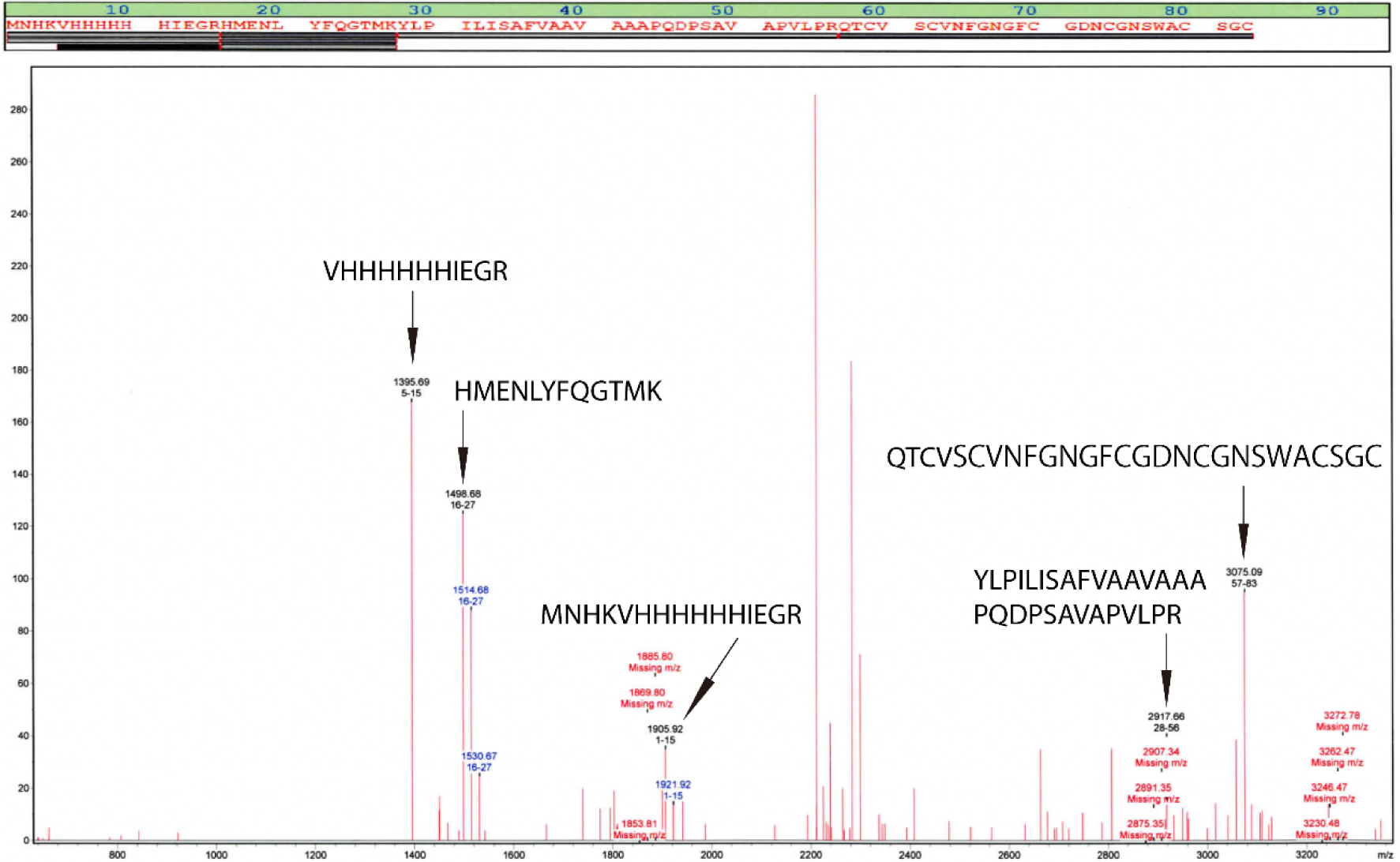
MALDI FT-ICR mass spectrometry of Cas1. The m/z signals in good agreement with the calculated mass are marked by the corresponding amino acid sequences. **<u>Related to</u> Figure 1 and Figure 1-table 1.**

**Figure 1-figure supplement 3.**
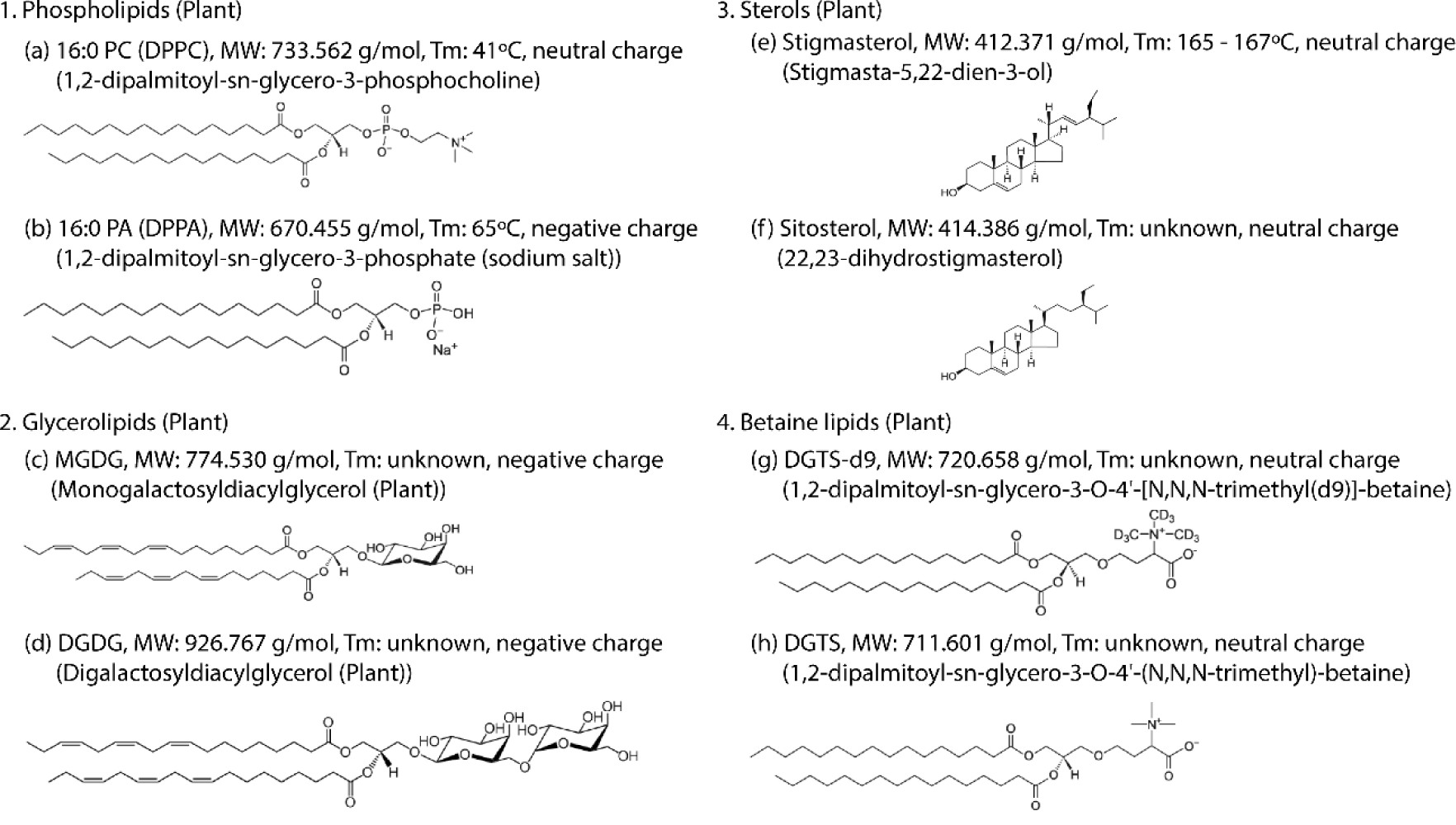
Structures and physical properties of different lipids available in cytoplasmic membranes of plant cells. Tm: melting temperature. Data were obtained from Avanti Polar Lipids (Alabama, USA). **<u>Related to</u> Figures 1<u>, 2, 3, and 4</u>**.

**Figure 6-figure supplement 1.**
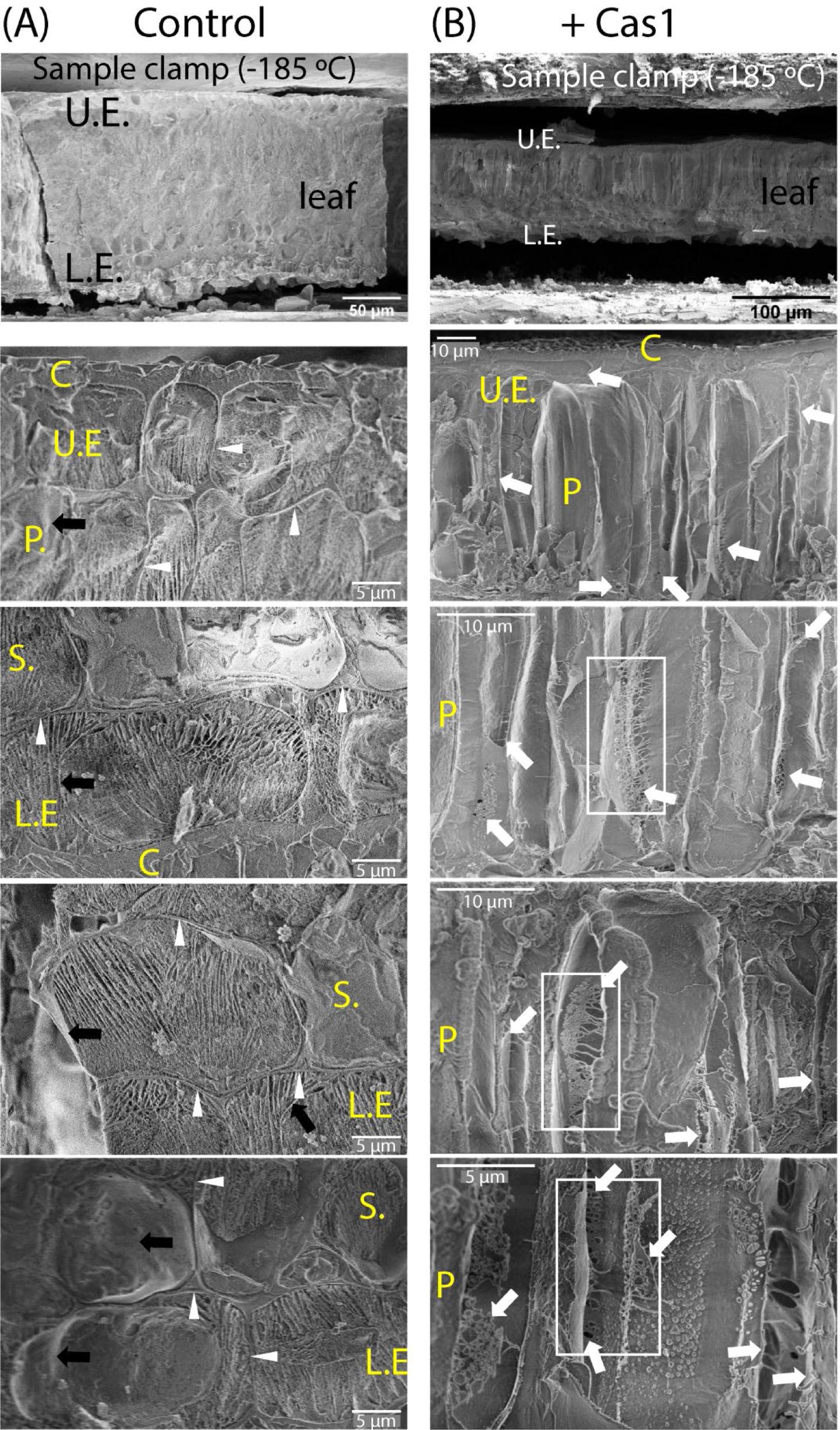
Cryo-SEM images of cellular and tissue structures of control and lesion spots in rubber leaves (rubber clone RRIV 4) caused by Cas1. Black and white arrows denote healthy and unhealthy cytoplasmic membranes, respectively. White arrowheads denote the cell wall. C: cuticle; U.E.: upper epidermis; L.E.: lower epidermis; P.: palisade; S.: spongy. Rectangles denote biofilm-like structures of Cas1 formed on cytoplasmic membrane. **<u>Related to</u> Figure 6**.

## Table supplements

**Figure 1-table 1.**
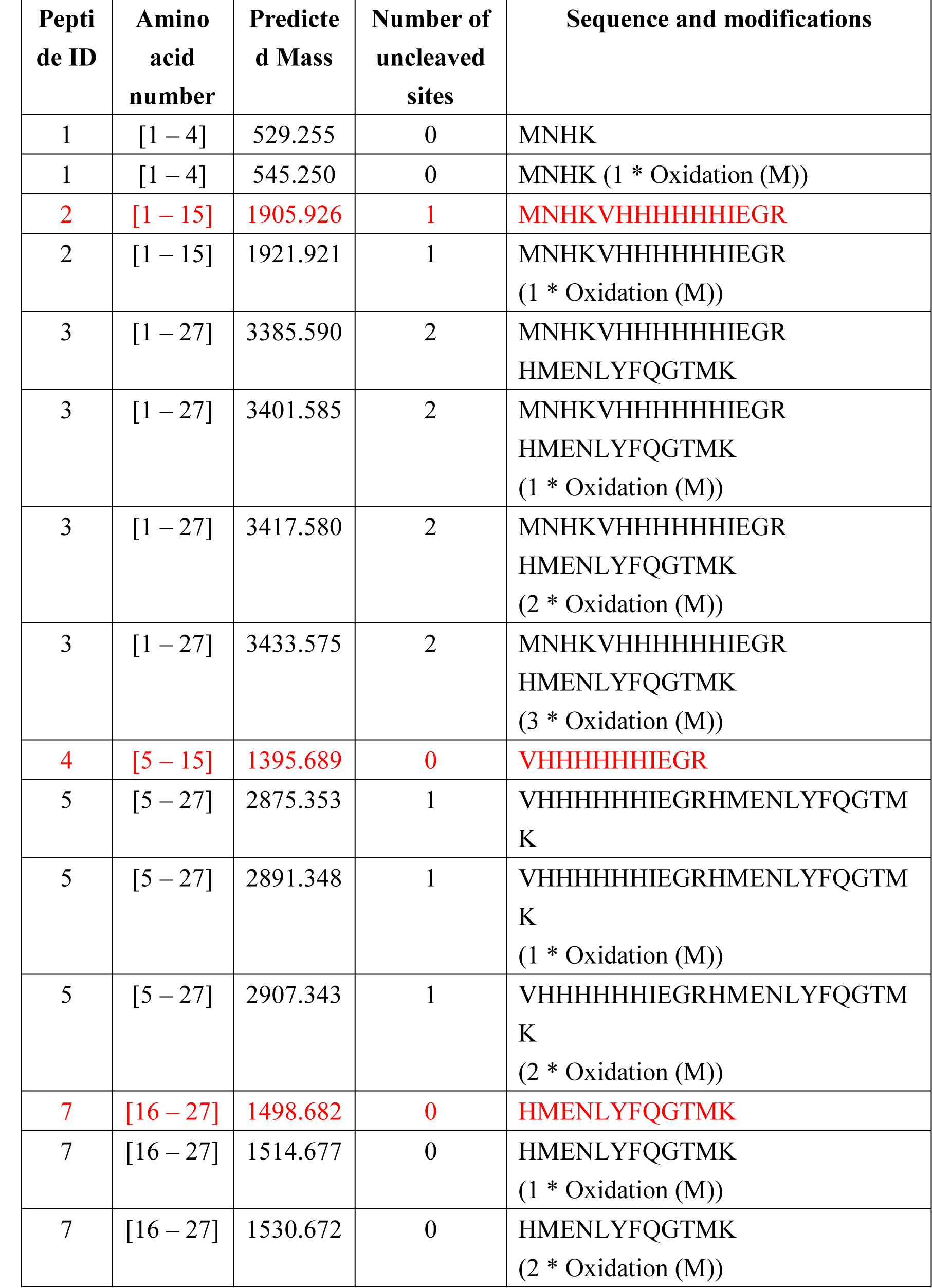
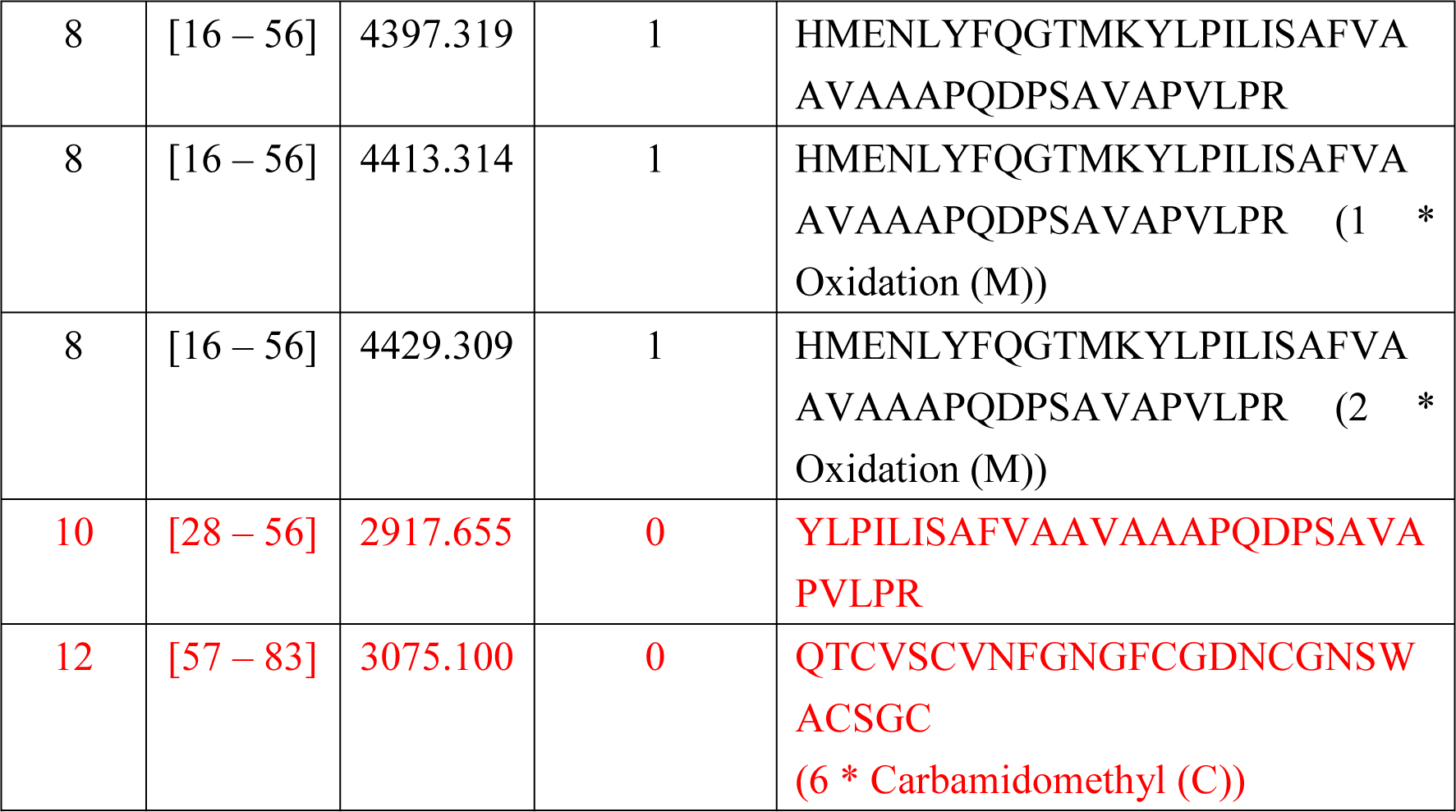
Peptides predicted for and detected (red characters) by MALDI FT-ICR mass spectrometry. **<u>Related to</u> Figure 1-figure supplement 2**

**Figure 1-table 2.**
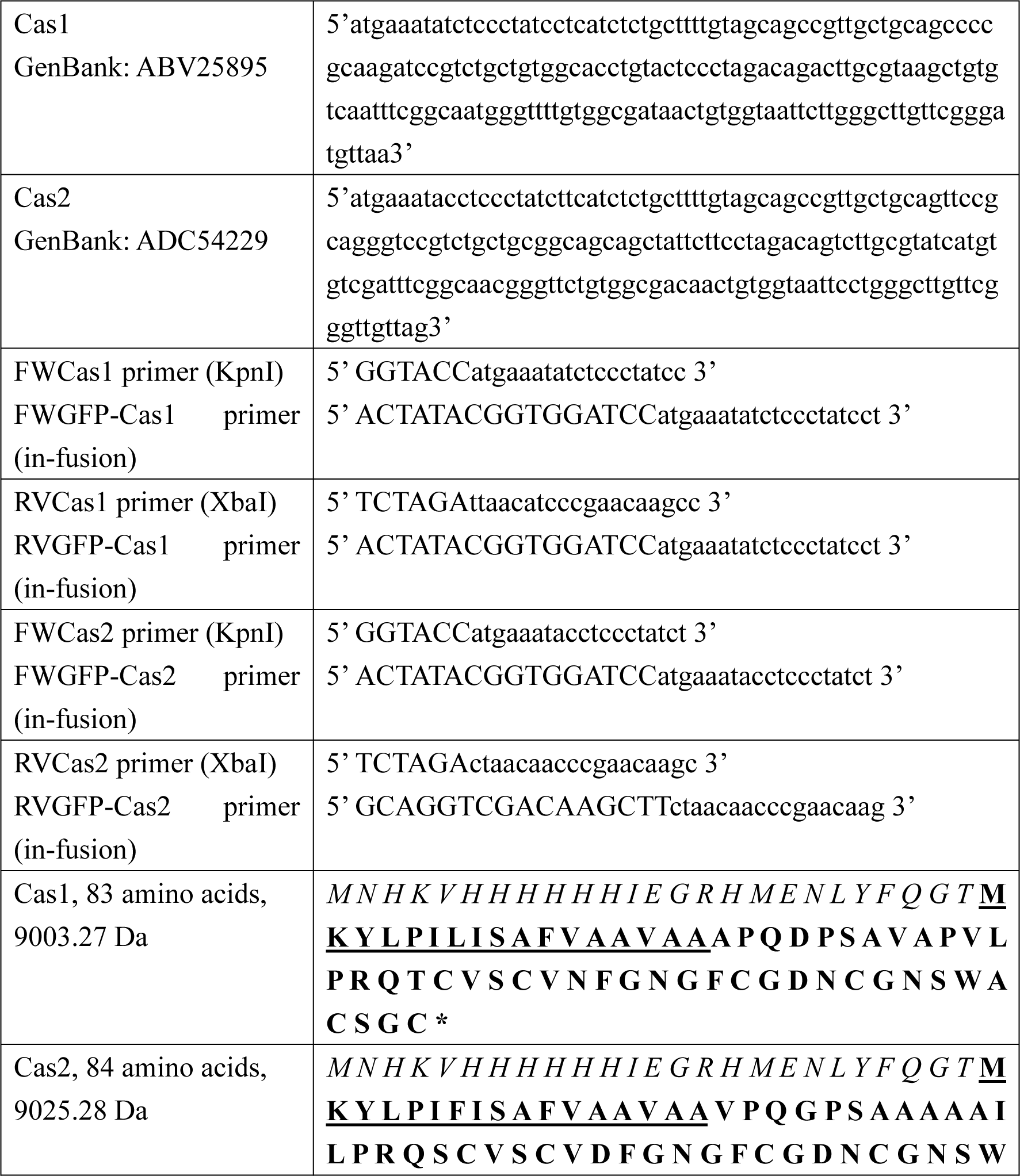
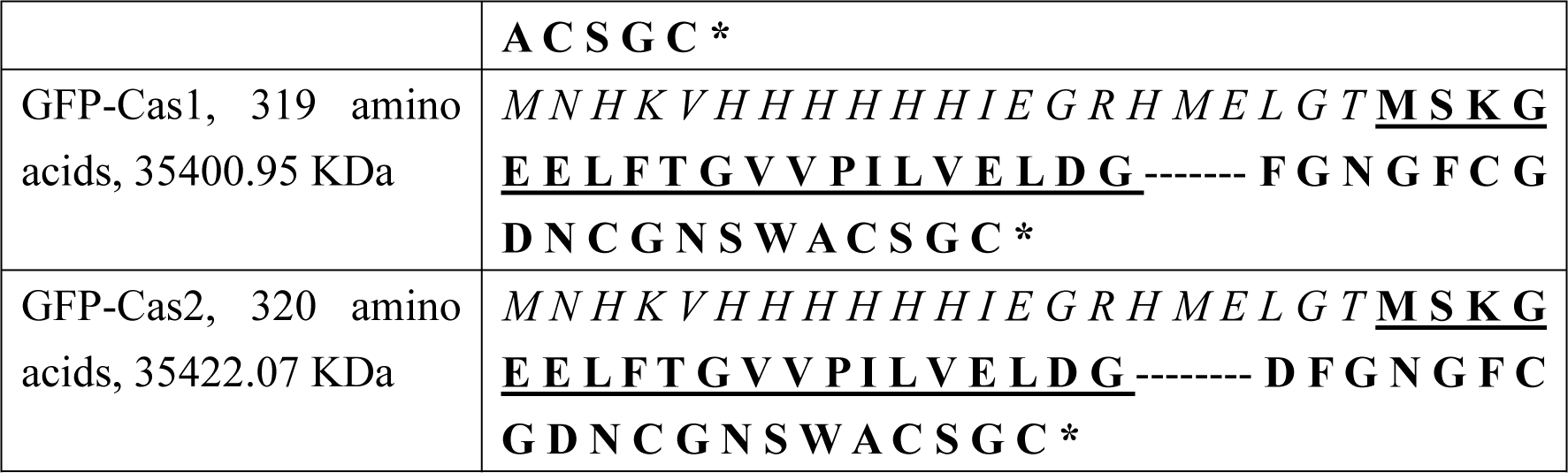
Information on oligonucleotide sequences of genes encoding Cas1 and Cas2 and amino acid sequences of recombinant Cas1, Cas2, GFP-Cas1, and GFP-Cas2 proteins. We explain amino acid sequences in Cas1 and Cas2; bold letters are full-length Cas1 and Cas2 proteins, in which the signal peptides are underlined letters followed by the putative mature domains. The italic letters indicate a 6xHis-tag followed by the amino acid linker on the N-terminal domain of Cas1 and Cas2 for purifying these recombinant proteins. For GFP-Cas1 and GFP-Cas2, bold letters are a fusion of GFP (underlined and bold letters) with Cas1 or Cas2 (bold letters). The italic letters indicate a 6xHis-tag followed by the amino acid linker. **<u>Related to</u> Figure 1-figure supplement 1.**

**Figure 4-table 1.**
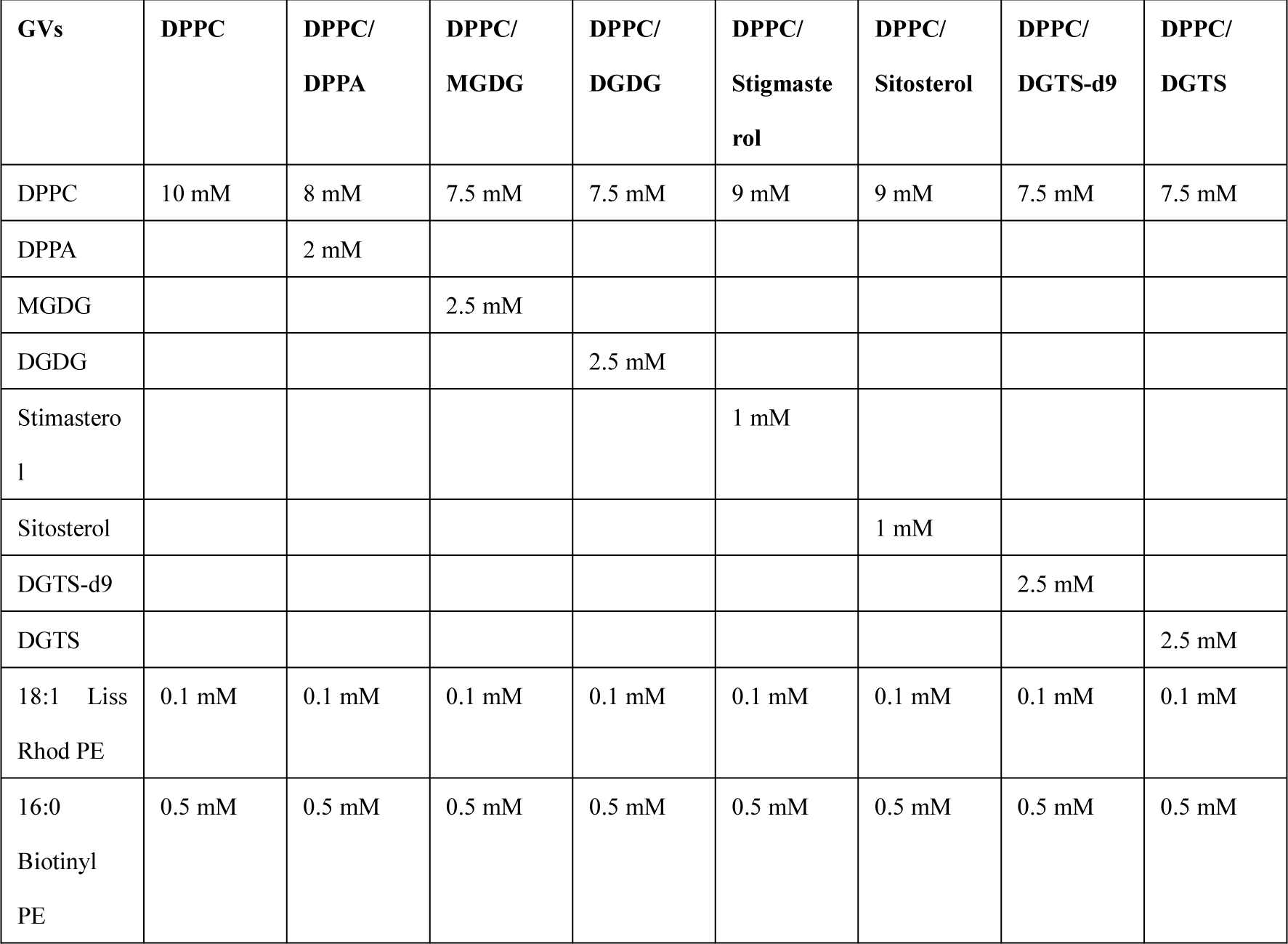
Lipid compositions for making composite giant liposome vesicles (GVs). Because the composite GVs contain both giant multilamellar vesicles (GMVs) and giant unilamellar vesicles (GUVs), we therefore use these composite giant vesicles as GVs. **<u>Related to</u> Figure 4**

## Video supplements

**Video 1**. Disruption of DPPC membrane lipids caused by Cas1. Briefly, DPPC lipid bilayers were formed on mica by rupturing DPPC liposomes. Then, 0.95 µM Cas1 was added, and HS-AFM imaging was carried out in C-buffer using BL-AC10DS-A2 cantilever deposited with a small Lab-made tip. AFM imaging rate: 2 fps. The Video plays at 30 fps. Bars: 50 nm. **<u>Related to Figure 1a1, Figure 1-figure supplement 3, and Table 1.</u>**

**Video 2**. Disruption of DPPC membrane lipids caused by Cas2. Briefly, DPPC lipid bilayers were formed on mica by rupturing DPPC liposomes. Then, 0.95 µM Cas2 was added, and HS-AFM imaging was carried out in C-buffer using BL-AC10DS-A2 cantilever deposited with a small Lab-made tip. AFM imaging rate: 2 fps. The Video plays at 30 fps. Bars: 50 nm. **<u>Related to Figure 1a2, Figure 1-figure supplement 3, and Table 1.</u>**

**Video 3**. Disruption of DPPC/DPPA membrane lipids caused by Cas1. Briefly, DPPC/DPPA (85/20 wt%) lipid bilayers were formed on mica by rupturing DPPC/DPPA liposomes. Then, 0.95 µM Cas1 was added, and HS-AFM imaging was carried out in C-buffer using BL-AC10DS-A2 cantilever deposited with a small Lab-made tip. AFM imaging rate: 2 fps. The Video plays at 30 fps. Bars: 50 nm. **<u>Related to Figure 1b1,</u> Figure 1<u>-figure supplement 3, and</u>** Table 1.

**Video 4**. Disruption of DPPC/DPPA membrane lipids caused by Cas2. Briefly, DPPC/DPPA (85/20 wt%) lipid bilayers were formed on mica by rupturing DPPC/DPPA liposomes. Then, 0.95 µM Cas2 was added, and HS-AFM imaging was carried out in C-buffer using BL-AC10DS-A2 cantilever deposited with a small Lab-made tip. AFM imaging rate: 2 fps. The Video plays at 30 fps. Bars: 50 nm. **<u>Related to Figure 1b2,</u> Figure 1<u>-figure supplement 3, and</u>** Table 1.

**Video 5**. Disruption of DPPC/MGDG membrane lipids caused by Cas1. Briefly, DPPC/MGDG (80/20 wt%) lipid bilayers were formed on mica by rupturing DPPC/MGDG liposomes. Then, 0.95 µM Cas1 was added, and HS-AFM imaging was carried out in C-buffer using BL-AC10DS-A2 cantilever deposited with a small Lab-made tip. AFM imaging rate: 2 fps. The Video plays at 30 fps. Bars: 50 nm. **<u>Related to Figure 1c1,</u> Figure 1<u>-figure supplement 3, and</u>** Table 1.

**Video 6**. Disruption of DPPC/MGDG membrane lipids caused by Cas2. Briefly, DPPC/MGDG (80/20 wt%) lipid bilayers were formed on mica by rupturing DPPC/MGDG liposomes. Then, 0.95 µM Cas2 was added, and HS-AFM imaging was carried out in C-buffer using BL-AC10DS-A2 cantilever deposited with a small Lab-made tip. AFM imaging rate: 2 fps. The Video plays at 30 fps. Bars: 50 nm. **<u>Related to</u>** <u>Figure 1c2,</u> Figure 1<u>-figure supplement 3, and</u> Table 1.

**Video 7**. Disruption of DPPC/DGDG membrane lipids caused by Cas1. Briefly, DPPC/DGDG (60/40 wt%) lipid bilayers were formed on mica by rupturing DPPC/DGDG liposomes. Then, 0.95 µM Cas1 was added, and HS-AFM imaging was carried out in C-buffer using BL-AC10DS-A2 cantilever deposited with a small Lab-made tip. AFM imaging rate: 2 fps. The Video plays at 30 fps. Bars: 50 nm. **<u>Related to Figure 1d1,</u> Figure 1<u>-figure supplement 3, and</u>** Table 1.

**Video 8**. Disruption of DPPC/DGDG membrane lipids caused by Cas2. Briefly, DPPC/DGDG (60/40 wt%) lipid bilayers were formed on mica by rupturing DPPC/DGDG liposomes. Then, 0.95 µM Cas2 was added, and HS-AFM imaging was carried out in C-buffer using BL-AC10DS-A2 cantilever deposited with a small Lab-made tip. AFM imaging rate: 2 fps. The Video plays at 30 fps. Bars: 50 nm. **<u>Related to Figure 1d2,</u> Figure 1<u>-figure supplement 3, and</u>** Table 1.

**Video 9**. Disruption of DPPC/stigmasterol membrane lipids caused by Cas1. Briefly, DPPC/stigmasterol (80/20 wt%) lipid bilayers were formed on mica by rupturing DPPC/stigmasterol liposomes. Then, 0.95 µM Cas1 was added, and HS-AFM imaging was carried out in C-buffer using BL-AC10DS-A2 cantilever deposited with a small Lab-made tip. AFM imaging rate: 2 fps. The Video plays at 30 fps. Bars: 50 nm. **<u>Related to Figure 1e1,</u> Figure 1<u>-figure supplement 3, and</u>** Table 1.

**Video 10**. Disruption of DPPC/stigmasterol membrane lipids caused by Cas2. Briefly, DPPC/stigmasterol (80/20 wt%) lipid bilayers were formed on mica by rupturing DPPC/stigmasterol liposomes. Then, 0.95 µM Cas2 was added, and HS-AFM imaging was carried out in C-buffer using BL-AC10DS-A2 cantilever deposited with a small Lab-made tip. AFM imaging rate: 2 fps. The Video plays at 30 fps. Bars: 50 nm. **<u>Related to Figure 1e2,</u> Figure 1<u>-figure supplement 3, and</u>** Table 1.

**Video 11**. Disruption of DPPC/sitosterol membrane lipids caused by Cas1. Briefly, DPPC/sitosterol (80/20 wt%) lipid bilayers were formed on mica by rupturing DPPC/Sitosterol liposomes. Then, 0.95 µM Cas1 was added, and HS-AFM imaging was carried out in C-buffer using BL-AC10DS-A2 cantilever deposited with a small Lab-made tip. AFM imaging rate: 2 fps. The Video plays at 30 fps. Bars: 50 nm. **<u>Related to Figure 1f1,</u> Figure 1<u>-figure supplement 3, and</u>** Table 1.

**Video 12**. Disruption of DPPC/sitosterol membrane lipids caused by Cas2. Briefly, DPPC/sitosterol (80/20 wt%) lipid bilayers were formed on mica by rupturing DPPC/Sitosterol liposomes. Then, 0.95 µM Cas2 was added, and HS-AFM imaging was carried out in C-buffer using BL-AC10DS-A2 cantilever deposited with a small Lab-made tip. AFM imaging rate: 2 fps. The Video plays at 30 fps. Bars: 50 nm. **<u>Related to</u>** <u>Figure 1f2,</u> Figure 1<u>-figure supplement 3, and</u> Table 1.

**Video 13**. Disruption of DPPC/DGTS-d9 membrane lipids caused by Cas1. Briefly, DPPC/DGTS-d9 (80/20 wt%) lipid bilayers were formed on mica by rupturing DPPC/DGTS-d9 liposomes. Then, 0.95 µM Cas1 was added, and HS-AFM imaging was carried out in C-buffer using BL-AC10DS-A2 cantilever deposited with a small Lab-made tip. AFM imaging rate: 2 fps. The Video plays at 30 fps. Bars: 50 nm. **<u>Related to Figure 1g1,</u> Figure 1<u>-figure supplement 3, and</u>** Table 1.

**Video 14**. Disruption of DPPC/DGTS-d9 membrane lipids caused by Cas2. Briefly, DPPC/DGTS-d9 (80/20 wt%) lipid bilayers were formed on mica by rupturing DPPC/DGTS-d9 liposomes. Then, 0.95 µM Cas2 was added, and HS-AFM imaging was carried out in C-buffer using BL-AC10DS-A2 cantilever deposited with a small Lab-made tip. AFM imaging rate: 2 fps. The Video plays at 30 fps. Bars: 50 nm. **<u>Related to Figure 1g2,</u> Figure 1<u>-figure supplement 3, and</u>** Table 1.

**Video 15**. Disruption of DPPC/DGTS membrane lipids caused by Cas1. Briefly, DPPC/DGTS (80/20 wt%) lipid bilayers were formed on mica by rupturing DPPC/DGTS liposomes. Then, 0.95 µM Cas1 was added, and HS-AFM imaging was carried out in C-buffer using BL-AC10DS-A2 cantilever deposited with a small Lab-made tip. AFM imaging rate: 2 fps. The Video plays at 30 fps. Bars: 50 nm. **<u>Related to Figure 1h1,</u> Figure 1<u>-figure supplement 3, and</u>** Table 1.

**Video 16**. Disruption of DPPC/DGTS membrane lipids caused by Cas2. Briefly, DPPC/DGTS (80/20 wt%) lipid bilayers were formed on mica by rupturing DPPC/DGTS liposomes. Then, 0.95 µM Cas2 was added, and HS-AFM imaging was carried out in C-buffer using BL-AC10DS-A2 cantilever deposited with a small Lab-made tip. AFM imaging rate: 2 fps. The Video plays at 30 fps. Bars: 50 nm. **<u>Related to Figure 1h2,</u> Figure 1<u>-figure supplement 3, and</u>** Table 1.

**Video 17**. Biofilm-like formation of Cas1/Cas2 mixtures onto DPPC/sitosterol membranes. Briefly, DPPC/sitosterol (80/20 wt%) lipid bilayers were formed on mica by rupturing DPPC/sitosterol liposomes. Then, 4.75 µM Cas1 and 4.75 µM Cas2 were added, and HS-AFM imaging was carried out in C-buffer using BL-AC10DS-A2 cantilever deposited with a small Lab-made tip. AFM imaging rate: 2 fps. The Video plays at 30 fps. Bars: 50 nm. **<u>Related to</u> Figure 3A.**

**Video 18**. Biofilm-like formation of Cas2 onto DPPC/sitosterol membranes. Briefly, DPPC/sitosterol (80/20 wt%) lipid bilayers were formed on mica by rupturing DPPC/sitosterol liposomes. Then, 4.75 µM Cas2 was added, and HS-AFM imaging was carried out in C-buffer using BL-AC10DS-A2 cantilever deposited with a small Lab-made tip. AFM imaging rate: 2 fps. The Video plays at 30 fps. Bars: 50 nm. **<u>Related to</u>** Figure 3B.

**Video 19**. Biofilm-like structures of Cas1/Cas2 mixtures were not formed onto DPPC/DGTS-d9 membranes. Briefly, DPPC/DGTS-d9 (80/20 wt%) lipid bilayers were formed on mica by rupturing DPPC/DGTS-d9 liposomes. Then, 4.75 µM Cas1 and 4.75 µM Cas2 were added, and HS-AFM imaging was carried out in C-buffer using BL- AC10DS-A2 cantilever deposited with a small Lab-made tip. AFM imaging rate: 2 fps. The Video plays at 30 fps. Bars: 50 nm. **<u>Related to</u> Figure 3C.**

